# Parallel genetic adaptation amidst a background of changing effective population sizes in divergent yellow perch (*Perca flavescens*) populations

**DOI:** 10.1101/2024.04.21.590447

**Authors:** Xiaoshen Yin, Claire E. Schraidt, Morgan M. Sparks, Peter T. Euclide, Tyler J. Hoyt, Carl R. Ruetz, Tomas O. Höök, Mark R. Christie

**Affiliations:** MOE Key Laboratory of Marine Genetics and Breeding, College of Marine Life Sciences, Ocean University of China, Qingdao 266003, China; Department of Biological Sciences, Purdue University; 915 W. State St., West Lafayette, Indiana 47907-2054 USA; Department of Forestry and Natural Resources, Purdue University; 715 W. State St., West Lafayette, Indiana 47907-2054 USA; Department of Ecology and Evolutionary Biology, Yale University; 165 Prospect Street, New Haven, Connecticut 06511 USA; US Forest Service, Rocky Mountain Research Station; 322 East Front Street, Boise, Idaho 83702 USA; Illinois-Indiana Sea Grant, Purdue University, West Lafayette, Indiana 47907 USA; Annis Water Resources Institute, Grand Valley State University; 740 W. Shoreline Dr., Muskegon, Michigan 49441 USA

## Abstract

Aquatic ecosystems are highly dynamic environments vulnerable to natural and anthropogenic disturbances. High-economic value fisheries are one of many ecosystem services affected by these disturbances and it is critical to accurately characterize the genetic diversity and effective population sizes of valuable fish stocks through time. We used genome-wide data to reconstruct the demographic histories of economically important yellow perch (*Perca flavescens*) populations. In two isolated and genetically divergent populations, we provide independent evidence for simultaneous increases in effective population sizes over both historic and contemporary time scales including negative genome-wide estimates of Tajima’s D, 3.1 times more SNPs than adjacent populations, and contemporary effective population sizes that have increased 10- and 47-fold from their minimum, respectively. The excess of segregating sites and negative Tajima’s D values likely arose from mutations accompanying historic population expansions with insufficient time for purifying selection, whereas linkage disequilibrium-based estimates of *Ne* also suggest contemporary increases that may have been driven by reduced fishing pressure or environmental remediation. We also identified parallel, genetic adaptation to reduced visual clarity in the same two habitats. These results suggest that the synchrony of key ecological and evolutionary processes can drive parallel demographic and evolutionary trajectories across independent populations.

## Introduction

Accurate estimates of effective population sizes (*Ne*) are critical for the successful management and conservation of exploited, threatened, and endangered species. The concept of *Ne* was first introduced in 1931, and often refers to the size of an idealized population with constant size and random mating that experiences an equivalent rate of genetic drift as the population under study (1). Consequently, estimates of effective population size can often be smaller than estimates of census population sizes (*Ne*/*N* ratio ranging from 10^-5^ to ∼0.5) (2–13). From a conservation and management perspective, effective population size matters in several key aspects: 1.) *Ne* is a major determinant of nucleotide variation (*i.e.*, *4Neμ* under a neutral model), 2.) *Ne* measures the expected rate of genetic drift, a major, stochastic, evolutionary force acting on genetic diversity across the genome and through generations, and 3.) *Ne* provides a unified frame of reference for comparisons across species or populations (14,15). Thus, changes in effective population sizes through time can provide information on how populations are responding to novel stressors, such as climate change (16–19), changes in food web dynamics (20), habit alteration and destruction (17,20), and overexploitation (16–18,21,22).

Accurate estimates of effective population sizes are particularly needed in fisheries to guide sustainable harvest practices in economically important populations. *Ne* is a critical metric for the successful assessment and management of fisheries stocks as it not only provides a fisheries-independent approach, avoiding the biases inherent in some field- or catch-based methods (23), but also reflects the adaptative potential of different stocks under constantly changing environments (23,24). Estimates of effective population sizes may forecast overharvest and potential collapses in commercial fisheries, as signaled by either substantial reductions in effective population sizes or increasing ratios of effective to census population sizes (*i.e.*, overharvest reduces both effective and census population sizes but reduces census population size more quickly, so the ratio of effective to census population size goes up) (4,25–27).

Nevertheless, several challenges remain for estimating effective population sizes in fish and fisheries. Estimates of *Ne* often include infinity when the signal for genetic variation arising from genetic drift cannot be estimated with high precision and is instead overwhelmed by noise arising from sampling error (28,29). Also, fish populations are often highly connected due to extensive larval dispersal and juvenile or adult migration (30–33) and such migration and subsequent gene flow can bias, or at the very least complicate estimates of *Ne* (14,34,35). Additional challenges include overlapping generations (34–36), varied genomic architectures (34), rare alleles (34), small sample sizes (34,35), sampling design (34), and sequencing or genotyping errors (34).

Despite these challenges, advances in high throughput sequencing and the influx of genomic data have resulted in the development of many methods (37–41) that more accurately infer temporal changes in effective population sizes for various data types, including polymorphic sites, short sequence blocks, and whole-genome data (14).

Empirical assessments of *Ne* over contemporary time scales (*e.g.*, decades to centuries) have largely demonstrated reductions in effective population sizes (16–22,42–45). Although rarely documented (6,45,46), contemporary increases in effective population sizes exist and can occur via one of three mechanisms. First, a reduction in the variance in reproductive success, for example by equalizing mating success throughout a single generation, can result in larger effective population sizes in the next generation (47,48), as demonstrated in captive breeding programs where equalization of full-sib family sizes is often recommended to increase effective population sizes (49–52). Second, introgression from neighboring, genetically divergent populations may generate biases in estimating *Ne*, the direction of which is determined by the magnitude and continuity of gene flow (34,35). Third, large, consistent increases in adult abundance within a population, perhaps due to improved environmental conditions and management policies, can also cause increases in effective population sizes (46). Over longer time scales, these last two mechanisms may also be accompanied by several other genetic indicators including negative values of Tajima’s D, shifts in site frequency spectra, and larger numbers of polymorphic loci with low minor allele frequency (14).

In this study, we explored changes in the effective population sizes and patterns of adaptive divergence in yellow perch (*Perca flavescens*) populations found throughout the main basin of Lake Michigan and two connected water bodies, Green Bay and Muskegon Lake (Fig. 1). Lake Michigan, with a drainage basin of about 118,000 km^2^, is the fourth largest freshwater lake in the world (53,54). Green Bay, an inlet of northwestern Lake Michigan, drains approximately 40,468 km^2^ while Muskegon Lake, a drowned river mouth (DRM) lake of 16.8 km^2^, receives water from the Muskegon River and drains directly into Lake Michigan. DRM lakes formed as the mouths of rivers were cut off from flowing into the Great Lakes following the retreat of glaciers. The melting river water was trapped behind land formations, carving out basins like Muskegon Lake, and eventually reconnecting after the buildup of water pressure created channels allowing them to once again flow into the Great Lakes (55,56). The main basin of Lake Michigan, Green Bay, and Muskegon Lake are all home to yellow perch, but are vastly different in both abiotic (*e.g.*, water clarity) and biotic (*e.g.*, community composition) characteristics and have distinct environmental histories. Yellow perch migrate between nearshore Lake Michigan and Muskegon Lake seasonally, but they do not appear to interbreed with resident populations (57,58). Previous work in this system using a RAD-Seq data set revealed that yellow perch in the main basin of Lake Michigan are genetically divergent from Green Bay (59,60), however the demographic and evolutionary histories of yellow perch from these three highly differentiated habitats have not yet been investigated (57–60).

**Fig. 1.**
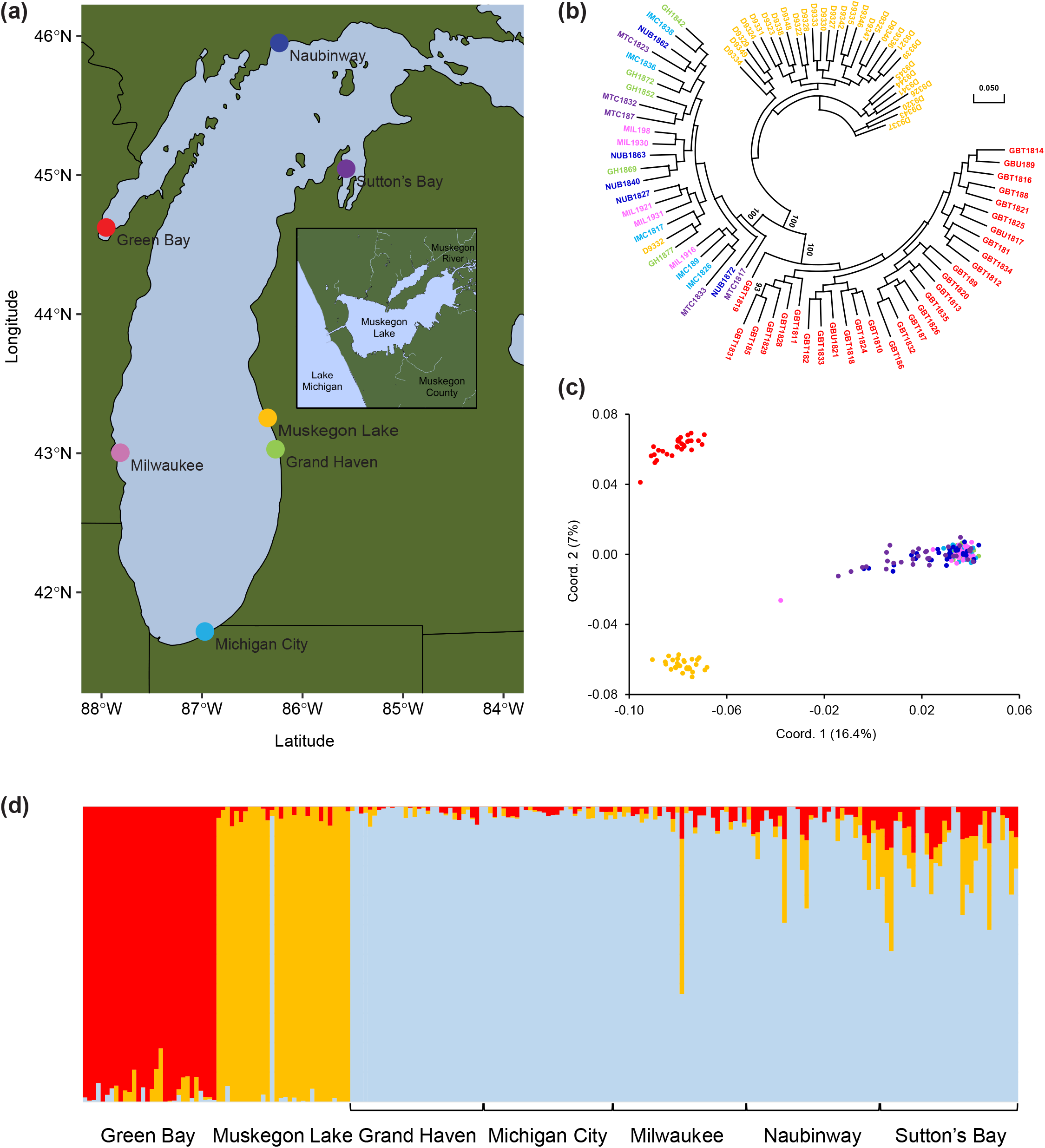
Population structure of Lake Michigan yellow perch populations. Yellow perch collected from seven Lake Michigan sites (a), including the comparatively isolated Muskegon Lake, which is connected to the main basin by a narrow channel (see inset). A phylogenetic tree illustrates three distinct groups: Green Bay, Muskegon Lake, and the five main basin sample sites (b). Twenty-five main basin individuals were randomly sampled for visual clarity; see Supplementary Fig. 1 for the full phylogenetic tree with all 210 individuals. Both principal coordinate analysis of 210 individuals (c) and admixture analyses (d) further support the delineation of samples into three discrete populations. Notice that one individual sampled in Muskegon Lake likely swam in from the main basin (Supplementary Fig. 2; see “Genetic divergence among yellow perch sample sites” in Supplementary Materials).

Yellow perch remain one of the most ecologically and economically important species throughout the Great Lakes. They are an abundant nearshore fish species, serve as important predators of small fishes and invertebrates, and are themselves important prey for larger fishes (see “Yellow perch life history” in Supplementary Materials for relevant life history information) (61). Historically, yellow perch supported commercially important fisheries throughout the Great Lakes region; in Lake Michigan alone, peak annual commercial harvest would now represent close to $16 million (US$) in dock-side value and much more at retail. However, Lake Michigan yellow perch populations began declining during the late 1980s and early 1990s, which led to closures of most yellow perch commercial fisheries. Here, we sequenced the complete genomes and reconstructed the demographic histories for 210 yellow perch collected from seven sample sites encompassing the main basin of Lake Michigan, Green Bay, and Muskegon Lake (Fig. 1a). Using genome-wide data, we ask the following questions: 1.) what are the differences in genetic differentiation and genetic diversity among sample sites?, 2.) what are the differences in *Ne* among sample sites using multiple approaches that can estimate both historical and more recent effective population sizes?, and 3.) what genes show signals of responding to selection in the three environmentally distinct regions (*i.e.*, main basin, Green Bay, and Muskegon Lake)? We identify three highly discrete populations, provide evidence for historic and contemporary increases in *Ne*, and find signatures of parallel, genetic adaptation.

## Methods

### Data collection

We sampled 210 adult yellow perch from seven locations (Green Bay, Muskegon Lake, Grand Haven, Michigan City, Milwaukee, Naubinway, Sutton’s Bay; Fig. 1a, Supplementary Table 1) in 2016, 2018 or 2019 using fyke netting, nighttime boat electrofishing, bottom trawls, gill netting and creel surveys, and conducted whole-genome resequencing to a depth of 12.50× after aligning to the yellow perch genome (62) (ranging from 5.93× to 18.58×; see “Whole-genome sequencing” in Supplementary Materials). We called SNPs using Genome Analysis Toolkit 4.1.9.0 (GATK 4.1.9.0) (63) (see “Variant calling and filtering” in Supplementary Materials).

We filtered SNPs by minor allele frequency (MAF) in two ways to create two data sets of SNPs suitable for different analyses (Supplementary Table 2): a **study-wide** data set and a **within-group** data set (see “Variant calling and filtering” in Supplementary Materials) (64). For population structure analyses, we used the study-wide data set, whereas we used the within-group data sets for all subsequent analyses (Supplementary Table 2).

### Population structure analysis

To infer the phylogenetic relationship among the seven sample sites, we constructed phylogenetic trees with 5,000 randomly selected SNPs using RAxML version 8.2.12 (65) and viewed the phylogenetic tree using MEGA11 (66) (see “Population structure analysis” in Supplementary Materials). To further investigate the population structure, we first conducted principal coordinate analysis on all individuals with 50,000 randomly selected SNPs in *hierfstat* in R 4.2.1 (67). Next, we conducted principal coordinate analysis with pairwise *FST* using GenAlEx 6.51b2 (68). Weir and Cockerham’s *FST* was calculated for each pair of sample sites using VCFtools (0.1.16) (69). Lastly, we estimated the ancestry for Green Bay, Muskegon Lake, and the five main basin sample sites, separately and jointly, with 10,000 randomly selected SNPs using Admixture 1.3 (70) by setting *K* from 1 to 10.

### Minor allele frequency, Tajima’s D, and genetic diversity

Using the within-group data sets, we calculated minor allele frequencies, Tajima’s D, and genetic diversity: 1) we calculated minor allele frequencies for seven sample sites and viewed its distribution by plotting density curves with unit bins of 0.015 with ggplot2 (71) and ggridges in R 4.2.1 (67), 2) we calculated Tajima’s D by setting the bin size to 5 MB with the within-group data set filtered by MAF ≥ 0.02 (Supplementary Table 2) using VCFtools (0.1.16), and 3) we estimated genetic diversity by calculating observed heterozygosity for every SNP (not including invariant sites) and averaged observed heterozygosity across overlapping windows of 5 MB using a step size of 2.5 MB across each chromosome. We also calculated nucleotide diversity (π) for each population with biallelic sites that are covered by at least three reads in each sample (including both polymorphic and invariant sites) using PIXY (72).

### Effective population size and demographic history

Using the within-group data sets (Supplementary Table 2), we estimated effective population size with 30,000 randomly selected SNPs (replicated 3 times per population) for each population using NeEstimator V2 (28) (see “Effective population size and demographic history” in Supplementary Materials). To further validate the *Ne* estimates, we estimated effective population size with 3,000 randomly selected SNPs from a previously obtained RAD-Seq data set (59), again replicated 3 times per sample site for each of 12 sample sites (see “Effective population size and demographic history” in Supplementary Materials), using the same options described above. Returning to our whole-genome data sets, we reconstructed demographic histories by running GONE (39) with all SNPs in the within-group data sets 100 times for each sample site. Specifically, we reconstructed the demographic history for Green Bay and Muskegon Lake by randomly selecting 30,000 SNPs from the within-group data set for each sample site (replicated 1000 times per sample site; Supplementary Table 2) and running GONE (39) for each of 1,000 randomly selected sets of 30,000 SNPs for each sample site using the default parameters. To examine changes in *Ne* over longer time scales, we also ran Stairway Plot 2 (38), for each sample site, with all biallelic SNPs at which genotype information is available for all samples in each of the seven sample sites, where generation time was estimated as four years +/- one year (see “Yellow perch life history” in Supplementary Materials).

### Candidate genes and gene ontology (GO) hierarchy networks underlying rapid adaptation

To identify genomic regions with overlapping windows of high *FST*, we first calculated pairwise Weir and Cockerham’s *FST* with separate VCF files for ten pairs of populations using VCFtools (0.1.16) (69) and identified outlier windows from a minimum of eight out of the ten pairwise population comparisons sharing at least one high *FST* SNP. We mapped SNPs on outlier windows to the yellow perch genome (62) to identify candidate outlier genes potentially explaining parallel genetic adaptation in Green Bay and Muskegon Lake in comparison to main basin populations. We annotated candidate outlier genes using UniProtKB ID via UniProt (https://www.uniprot.org/), with which we retrieved gene ontology (GO) terms for these outlier genes from Gene Ontology and GO Annotations (https://www.ebi.ac.uk/QuickGO/). Lastly, we constructed GO hierarchy networks of GO terms associated with at least two outlier genes in the domain of “biological process” for each pairwise comparison (*i.e.*, Green Bay or Muskegon Lake *vs.* main basin populations) using *metacoder* (73).

### Water clarity, water quality, and phenotypic measurements

Because many of the outlier genes identified from the previous analysis were related to vision, we investigated differences in water clarity, water quality, and yellow perch eye size between the main basin, Green Bay, and Muskegon Lake sample sites. Differences in eye size among those sample sites was not an *a priori* hypothesis but was rather an *a posteriori* hypothesis guided by our candidate genes (see Discussion). Water clarity was estimated using Secchi depth readings collected between June 1 and August 30 from Green Bay (320 readings), Muskegon Lake (72 readings), and the southern basin of Lake Michigan (180 measurements) between 2010 and 2022 (see “Water clarity, water quality, and phenotypic measurements” in Supplementary Materials). We conducted an Analysis of Variance (ANOVA) to test differences in Secchi depths among Green Bay, Muskegon Lake, and the southern basin of Lake Michigan.

To determine whether eye diameter differed among Green Bay, Muskegon Lake, and Lake Michigan, we collected yellow perch from the three locations, measured their eye diameter, and analyzed yellow perch with total lengths of 150-250 mm to ensure comparisons of yellow perch with similar sizes among the three locations (Green Bay: *n=31*, Muskegon Lake: *n=39*, Grand Haven: *n=27*; see “Water clarity, water quality, and phenotypic measurements” in Supplementary Materials). With this subset of fish, we used Analysis of Covariance (ANCOVA) to test whether eye diameter differed among the three locations while accounting for total length as a covariate (including interaction between location and covariate). We compared least-squares means with Tukey’s method using the “emmeans” function in the *emmeans* package in R 4.2.1 (67).

## Results

### Population structure

We mapped an average of 99.6% of 15.8 billion, 150 bp paired-end reads to the yellow perch genome (62) and, on average, over 90% and 83% of reads had a mapping quality equal to or greater than 20 or 60, respectively (Supplementary Table 3). The study-wide data set, which was used for population genetic and stock identification analyses, resulted in 714,666 SNPs per sample site while the within-group data set, which was used for demographic analyses, resulted in an average of 1,160,504 SNPs with less than 0.9% of SNPs out of Hardy-Weinberg equilibrium in each sample site (Supplementary Table 2, Supplementary Table 4). After inferring phylogenetic relationships among populations, we found that Green Bay, Muskegon Lake, and the main basin sample sites (*i.e.*, Grand Haven, Michigan City, Milwaukee, Naubinway, and Sutton’s Bay) form three distinct lineages (Fig. 1b, Supplementary Fig. 1). By conducting principal coordinate analysis with all individuals, we found further support that Green Bay, Muskegon Lake, and the main basin sample sites are clearly separated from each other (Fig. 1c, Supplementary Fig. 2a,b,c), except for one Muskegon Lake sample clustering with main basin populations and one Milwaukee main basin sample falling comparatively further away from the main basin populations (Fig. 1c, Supplementary Fig. 2d,e,f; see “Yellow perch life history” and “Genetic divergence among yellow perch sample sites” in Supplementary Materials). This clear delineation of three populations was further supported by principal coordinate analysis on the mean pairwise estimates of *FST* where we detected high genetic differentiation between Green Bay and Muskegon Lake (*FST* = 0.0520), both of which are genetically divergent from the five main basin sample sites (*FST* = 0.0550 to 0.0667, 0.0460 to 0.0592, respectively; Supplementary Table 5, Supplementary Fig. 3a). By contrast, the five main basin sample sites are comparatively similar to each other, with mean pairwise *FST* ranging from −0.0002 to 0.0041 (Supplementary Table 5, Supplementary Fig. 3b). The optimal number of clusters/populations identified with admixture analysis (70) for all populations was 3: Green Bay, Muskegon Lake, and the main basin sample sites (Fig. 1d, Supplementary Fig. 4).

### Numbers of SNPs, minor allele frequency, and Tajima’s D

When using the within-group data set, we detected 2.8-3.4 times more polymorphic SNPs in Green Bay and Muskegon Lake sample sites, respectively, in comparison to the main basin sample sites (2,186,768 and 2,275,935 *vs.* 670,578 to 776,088 SNPs in main basin populations; Fig. 2a). The median minor allele frequency of SNPs was 0.086 and 0.083 in Green Bay and Muskegon Lake, respectively, which was 0.46 times lower than that in main basin populations, which ranged from 0.183 to 0.185 (Fig. 2b). These differences in the number of SNPs and median minor allele frequencies were not driven by differences in sample sizes (Supplementary Table 1), sequencing quality (see “Whole-genome sequencing” in Supplementary Materials), sequencing depth (Supplementary Table 6), read depth of aligned reads (Supplementary Table 7), or missing data (see “Variant calling and filtering” in Supplementary Materials). Using the within-group data set, we observed consistently positive Tajima’s D values in the main basin populations, varying from 0.332 to 0.428 on average, which is suggestive of slight population declines in the main basin (Fig. 2c). By contrast, Green Bay and Muskegon Lake populations had a genome-wide average Tajima’s D of −0.632 and −0.603, respectively (Fig. 2c), which is suggestive of a recent (over evolutionary time scales) demographic expansion.

**Fig. 2.**
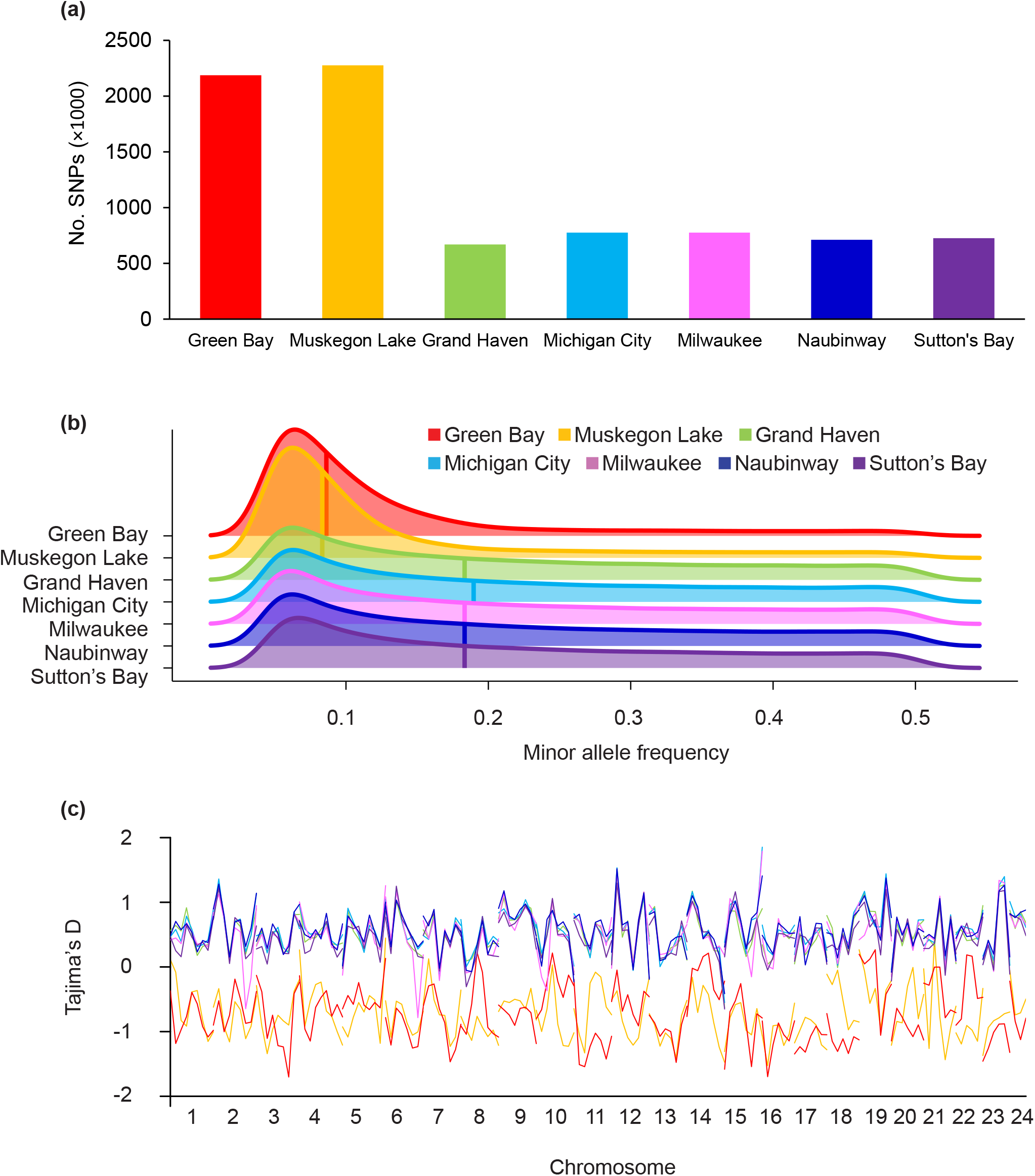
Numbers of single nucleotide polymorphisms (SNPs), minor allele frequency, and Tajima’s D. Green Bay and Muskegon Lake populations have 2.8-3.4 times more SNPs than the five main basin sample sites (a), despite all sample sites having equal sample sizes, sequencing quality, read depth, and missing data. Both Green Bay and Muskegon Lake also have a lower median minor allele frequency compared to the five main basin populations (b) here illustrated with a density plot where the median minor allele frequency is depicted with a solid vertical line. The negative Tajima’s D for both Green Bay and Muskegon Lake populations suggests recent population expansion (c). Collectively, these results suggest that the three populations (Green Bay, Muskegon Lake, and the main basin sample sites) have different demographic histories.

### Effective population size, genetic diversity, and demographic history

In comparison to the main basin sample sites, Green Bay and Muskegon Lake have significantly lower effective population sizes; the point estimates using whole-genome sequencing data for Green Bay and Muskegon Lake were 23 and 90, respectively, by NeEstimator V2 and 46 and 159, respectively, by GONE (Fig. 3a; Supplementary Table 8). Effective population size estimates by NeEstimator V2 using our RAD-Seq data set with larger numbers of samples (59), but smaller numbers of loci, suggest that southern and middle Green Bay (SGB, MEN) have smaller effective population sizes than northern Green Bay (BDN, LBD), with high levels of concurrence between both data sets, suggesting that main basin sample sites have considerably larger effective population sizes (Fig. 3b). Genetic diversity of main basin sample sites, with a mean observed heterozygosity ranging from 0.312 to 0.323, was considerably higher than that of Green Bay and Muskegon Lake populations, with a mean observed heterozygosity of 0.208 and 0.206, respectively (Fig. 3c, Supplementary Fig. 5). It should be noted that we calculated genetic diversity (*i.e.*, observed heterozygosity) using only polymorphic sites found in each within-group data set, meaning that the sets of SNPs for calculation differ among populations. Therefore, despite Green Bay and Muskegon Lake having a greater number of SNPs, the lower observed heterozygosity is predicted from theory given the much lower average minor allele frequencies. By contrast, estimates of nucleotide diversity (π), which is calculated using both polymorphic and invariant sites, were nearly three times higher in Green Bay and Muskegon Lake in comparison to the Lake Michigan main basin samples (Supplementary Fig. 6). Using GONE, we reconstructed contemporary demographic histories and found that Green Bay and Muskegon Lake experienced rapid growth of *Ne* over the last 10 to 30 generations (Fig. 3d,e, Supplementary Fig. 7). Stairway Plot 2 (38), based on site frequency spectra, suggests that increases in *Ne* in Green Bay and Muskegon Lake also occurred around 2000 years ago and even longer for main basin populations (Supplementary Fig. 8, Supplementary Fig. 9, Supplementary Fig. 10).

**Fig. 3.**
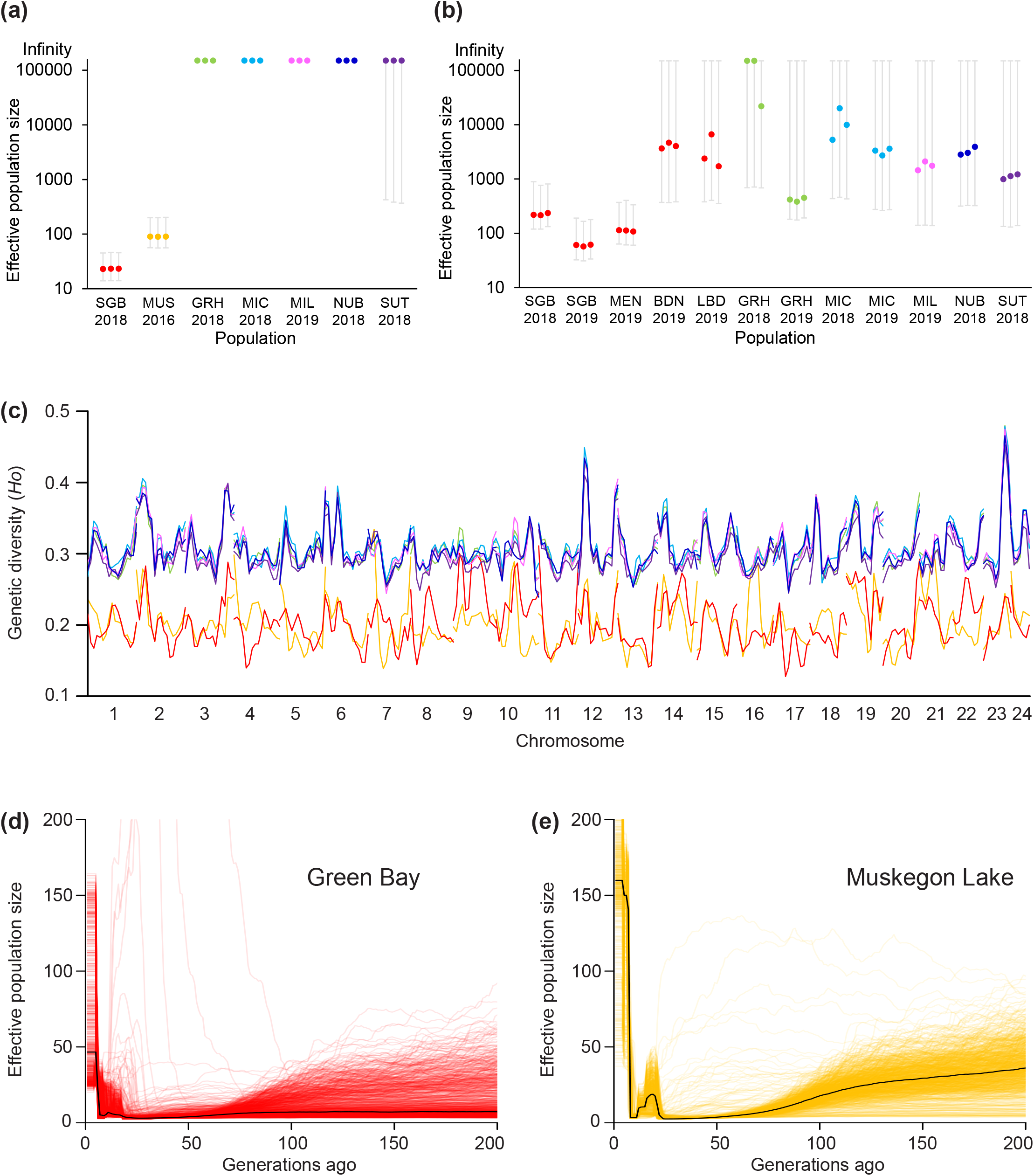
Demographic history of Lake Michigan yellow perch populations. Green Bay and Muskegon Lake sample sites have substantially lower effective population sizes, which average 23 and 90 estimated by NeEstimator V2, respectively, in comparison to main basin populations (a, estimates based on whole-genome sequencing data obtained in this study; b, estimates based on RAD-Seq data obtained from Schraidt *et al*.; note that y-axis is on a base 10 logarithmic scale; SGB: South Green Bay, MUS: Muskegon Lake, GRH: Grand Haven, MIC: Michigan City, MIL: Milwaukee, NUB: Naubinway, SUT: Sutton’s Bay, MEN: Menominee, BDN: Big Bay de Noc, LBD: Little Bay de Noc). Green Bay and Muskegon Lake have lower genetic diversity (c) than main basin populations, measured here as observed heterozygosity averaged across overlapping windows of 5 MB using a step size of 2.5 MB across each chromosome. Notice that the main basin heterozygosity estimates are highly correlated, confirming higher gene flow among main basin sample sites. The demographic history, as reconstructed by GONE, demonstrates a rapid increase in effective population size in Green Bay and Muskegon Lake since 10 to 30 generations ago (d,e).

### Candidate genes underlying genetic adaptation

Given the high levels of genetic divergence between Green Bay, Muskegon Lake, and the main basin sites, we used a genome-wide scan to determine if any regions of the genome are suggestive of genetic adaptation to the different environments (74,75). We retained a total of 13 outlier windows from nine chromosomes that were shared in at least 80% of comparisons between Green Bay and the main basin collection sites as well as Muskegon Lake and the main basin populations (see Methods; Supplementary Table 9). The 13 outlier windows contained an average of 1,427 SNPS (range: 363-3,203) and an average of 225 high *FST* SNPs (*FST* ≥ 0.6; range 82-360 SNPs per window) (Fig. 4a). SNPs within these 13 outlier windows mapped to 36 genes, among which three were located within a single outlier window on chromosome 4 (chromosome 4: 34762415-34852528). The three genes, *opn1sw*, *opn1lw*, and *opn1lw2*, encode visual pigments, light-absorbing molecules mediating vision. Out of the three visual pigment- encoding genes, *opn1lw2* has two outlier SNPs with *FST* ≥ 0.5 in comparisons between Green Bay and the main basin sample sites, and the exact same two outlier SNPs, one with *FST* ≥ 0.76 and the other with *FST* ≥ 0.46, were found in comparisons between Muskegon Lake and the main basin sample sites (Fig. 4a). The observation that three visual pigment-encoding genes were found on the same outlier window in all ten pairwise comparisons suggests that this region has responded to recent positive and parallel selection. Across all outlier windows, seven major categories of biological processes were detected between Green Bay and the main basin sample sites and Muskegon Lake and the main basin sample sites: multicellular organismal process, response to stimulus, biological regulation, cellular process, developmental process, localization, and metabolic process (Fig. 4b). Multiple visual perception-related biological processes, including detection of visible light, absorption of visible light, visual perception, cellular response to light stimulus, and phototransduction, provide further support for parallel adaptive differences in vision in Green Bay and Muskegon Lake yellow perch populations (Fig. 4b).

**Fig. 4.**
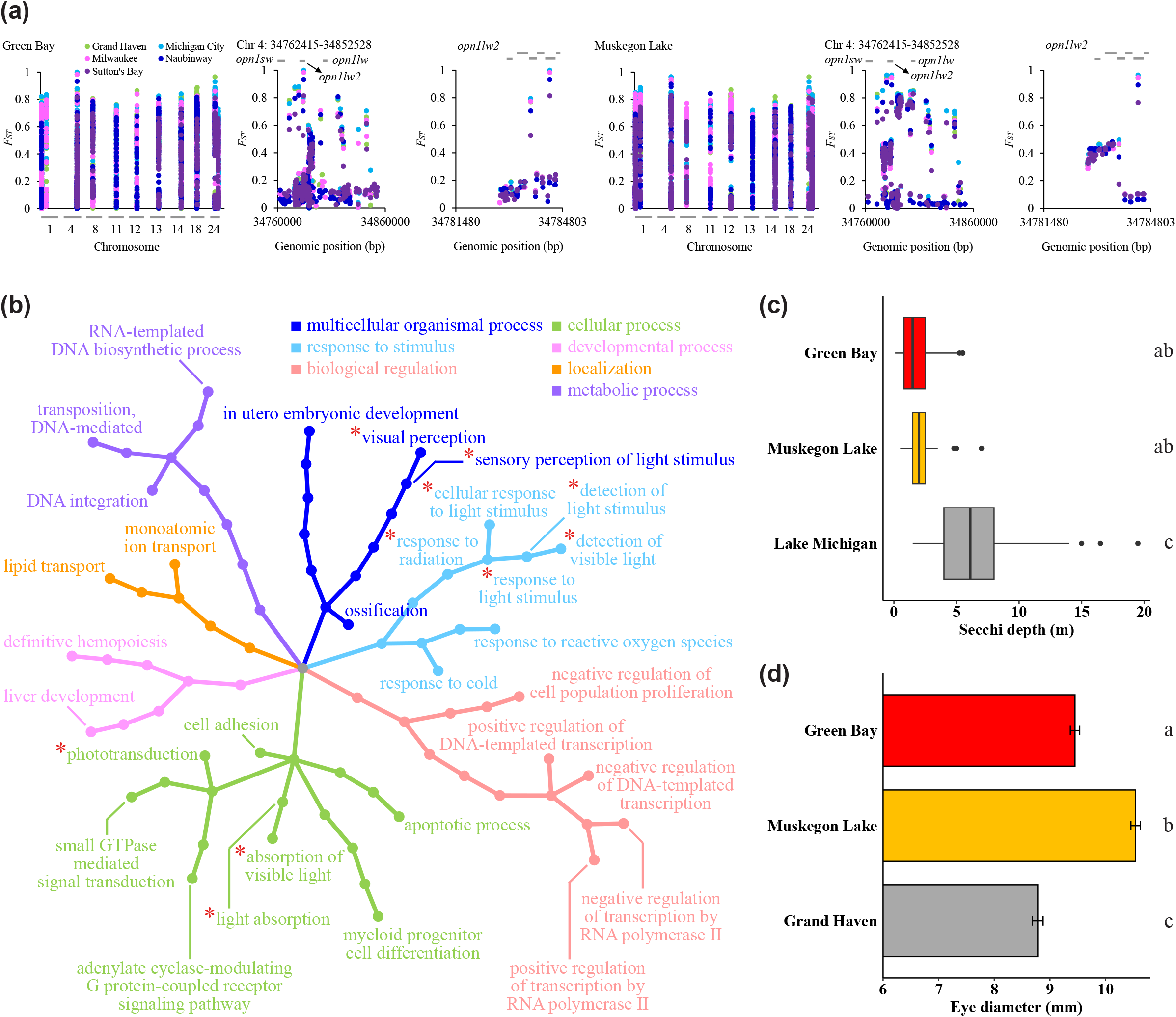
Candidate genes, biological processes, and phenotypes involved in parallel genetic adaptation. A total of 13 outlier windows were detected in comparisons between both Green Bay and the main basin sample sites and Muskegon Lake and the main basin sample sites (a). The outlier window on chromosome 4 contains visual pigment encoding genes, such as *opn1lw2*. The coding regions, as indicated by horizontal gray bars, of these outlier genes contain SNPs that are highly differentiated (a). Seven major categories of biological processes are potentially involved in adaptive differentiation in Green Bay and Muskegon Lake in comparison to main basin sample sites, among which visual perception-related processes (highlighted with red asterisks) further support parallel adaptive differences in vision in Green Bay and Muskegon Lake (b). The genetic differentiation at *opn1lw2* is likely associated with water clarity, as evidenced by significant differences in Secchi depths (c) and different eye sizes (d) between Green Bay or Muskegon Lake and main basin populations (*i.e.*, Lake Michigan, Grand Haven). The Secchi depth in Green Bay and Muskegon Lake is significantly smaller than that in the southern basin of Lake Michigan (*P* < 0.001; c) and least-squares means of yellow perch eye diameter in Green Bay and Muskegon Lake is significantly larger than that in Grand Haven (*P* < 0.001; d). We accounted for differences in total length when comparing eye diameter of yellow perch collected from Green Bay, Muskegon Lake, and Grand Haven (Supplementary Fig. 11; see Methods).

Using these candidate loci, we hypothesized that water clarity and eye size would differ between the two sites (Green Bay and Muskegon Lake) and the main basin. We found significant differences in water clarity between the main basin of Lake Michigan (low turbidity, high visual clarity; mean Secchi depth ± s.d.: 6.10 ± 3.23 m) and both Green Bay (high turbidity, low visual clarity; 1.59 ± 1.01 m) and Muskegon Lake (high turbidity, low visual clarity; 2.48 ± 1.23 m; ANOVA, *P* < 0.001; Fig. 4c). We also found that yellow perch eye diameter differed among locations (ANCOVA, *F*2,91 = 12.03; *P* < 0.001) and significantly increased with total length (ANCOVA, *F*1,91 = 45.67; *P* < 0.001), but the interaction was not significant (ANCOVA, *F*2,91 = 0.11; *P* = 0.779; Supplementary Fig. 11). Yellow perch eye diameter was significantly larger for Muskegon Lake yellow perch in comparison to fish collected from the main basin (Grand Haven; *P* < 0.001) and Green Bay (near Menominee; *P* < 0.001). Eye diameter for yellow perch from Green Bay was also significantly larger than fish from the main basin (Grand Haven; *P* < 0.001; Fig. 4d).

## Discussion

Using whole-genome sequence data from 210 yellow perch, we observed a clear delineation of three populations and multiple genomic signatures associated with historic and contemporary increases in *Ne* in Green Bay and Muskegon Lake where: (1) effective population sizes rapidly increased over recent generations, (2) Tajima’s D estimates were negative across the entire genome, and (3) a factor of 2.8-3.4 more polymorphic SNPs were found in Green Bay and Muskegon Lake than the main basin collection sites, where the majority of these additional SNPs occur at low frequency. The additional SNPs likely come from an excess of mutations that accumulated during population expansions that occurred over historic time scales (*i.e.*, thousands of years). This last observation is consistent with a large population expansion over historic time scales (*e.g.*, over thousands of years), as indicated by Stairway Plot 2 (Supplementary Fig. 9), where there has not been sufficient time for purifying selection to remove deleterious alleles (76). Given the substantial water quality and habitat improvements in Green Bay and Muskegon Lake driven by environmental remediation initiatives implemented in the early 1970s (Supplementary Fig. 12, Supplementary Fig. 13; see “Supplementary Discussion” in Supplementary Materials), we should not have been surprised to see an increase in *Ne*. Nevertheless, empirical documentation of contemporary increases in *Ne* remains rare and this result was unexpected.

To what time period do our demographic inferences apply? Linkage disequilibrium based methods of *Ne*, such as those depicted in Fig. 3, provide recent and contemporary estimates of effective population size (77). We used the program GONE to estimate recent changes to *Ne* over time and the results suggest that there have been contemporary increases in *Ne* in both Green Bay and Muskegon Lake. However, the exact date for when these increases in *Ne* began remains imprecise. A recent study with pink salmon (*Oncorhynchus gorbuscha*) and a known date of an extreme founder event illustrated that GONE may have missed the timing of the event by as much as six generations (45). Nevertheless, the known abundance patterns in salmon and other species are well characterized with this approach (45,78) and our results clearly illustrate that effective population sizes have been increasing in both Muskegon Lake and Green Bay in the last 10 to 30 generations, with larger increases occurring in Muskegon Lake. Our inferences based on Tajima’s D, the number of segregating loci, and site frequency spectra suggest much larger increases in *Ne* that occurred over historic time scales (centuries, not decades) (Fig. 2; Supplementary Fig. 9). This result contrasts with those from main basin sites, which all show increases in *Ne* starting more than 3000 years ago and no signs of a historic bottleneck (Supplementary Fig. 9). We speculate that both the Green Bay and the Muskegon Lake yellow perch populations were founded much more recently than the main basin populations or that both went through a large, unknown, bottleneck event approximately 2,000 years ago.

We also found evidence of parallel evolution in response to reduced water clarity in both Green Bay and Muskegon Lake yellow perch. In comparison to the main basin of Lake Michigan, both Green Bay and Muskegon Lake exhibit high turbidity and low visual clarity, as indicated by shallower Secchi depths. Such differences in water clarity and ambient light conditions could have driven the significant allele frequency shifts in the coding sequences of photopigment encoding genes and triggered a set of biological processes contributing to light detection and absorption, visual perception, and phototransduction, a result that implies parallel genetic adaptation in Green Bay and Muskegon Lake yellow perch in response to local environmental conditions. Differences in vision and eye size, while accounting for allometry, among those sample sites was not an *a priori* hypothesis, although it could have been had we thought about the differences in water clarity among the sample sites. Nevertheless, using whole-genome sequencing to guide phenotypic and evolutionary predictions does suggest a promising avenue for the future of genomic tools (79). Future studies should investigate linkages between candidate genes and their patterns of expression in addition to the structural and functional analyses of the proteins they encode. Additional phenotyping efforts and comparisons among multiple drowned river mouth lakes would also be useful. The observation that both genetic and phenotypic changes to reduced water clarity occurred in genetically divergent populations suggests that a match between vision-related traits and environmental characteristics is an important component of fitness in yellow perch populations (80); environmental changes that cause a mismatch may result in declines in abundance.

Several caveats are worth discussing. First, linkage disequilibrium-based estimates of effective population sizes may not accurately estimate effective population sizes prior to bottleneck events (45,81). As such, the effective population sizes of yellow perch in Green Bay and Muskegon Lake prior to around 30 generations ago were likely much larger than present day estimates but cannot be estimated using those approaches. Also, contemporary effective population size may be underestimated by linkage disequilibrium-based methods due to overlapping generations of yellow perch and the mixed-age adult samples under study (82).

Demographic estimates by GONE may also be unreliable for the most recent several generations. However, despite the wide range of estimates for recent generations, the general trend still indicates an increase in effective population size that appears to have persisted in Green Bay and Muskegon Lake for the past 10 to 30 generations. Perhaps most importantly, some of our demographic results could also be explained by gene flow from divergent populations (83,84); however, we do not think this to be the case for three reasons. First, PCA, phylogenetic, and Admixture analyses support a clear separation of Green Bay, Muskegon Lake, and Lake Michigan main basin populations and suggest the absence of admixed individuals in both Green Bay and Muskegon Lake. Even if all the main basin loci were introduced into Muskegon Lake samples via gene flow (which is unlikely), this mechanism could only account for 24% of the 2,275,935 SNPs found in Muskegon Lake (*i.e.*, 535,860 SNPs shared by Muskegon Lake and Grand Haven divided by 2,275,935 Muskegon Lake SNPs; Fig. 2a). Second, over shorter time scales, estimates of contemporary admixture within Green Bay and Muskegon Lake samples suggest a single, randomly mating population (as do the small number of loci out of Hardy-Weinberg equilibrium) (Supplementary Fig. 4; Supplementary Table 4). Third, over longer time scales, the high rates of introgression required to explain these results would also be likely to swamp any signal of parallel genetic adaptation (Fig. 4) and would also leave more time for purifying selection to remove deleterious loci. Note that other factors that can increase *Ne* through time (*e.g.*, decreased variance in reproductive success) are unlikely to be accompanied by the other genomic patterns we report (*i.e.*, negative Tajima’s D, larger numbers of SNPs).

Thus, we think the most parsimonious explanation for our results may be large, simultaneous increases in population sizes in Green Bay and Muskegon Lake yellow perch. Interestingly, fisheries independent data suggest that yellow perch have been declining recently in the main basin (85) and the positive, albeit small, genome-wide estimates of Tajima’s D suggest that these populations should be closely monitored in the coming decades. While we are aware that correlation does not equal causation, we speculate that the recent increases in *Ne* in Green Bay and Muskegon Lake could be a result of substantial water quality and habitat improvements driven by a set of environmental remediation initiatives implemented in the early 1970s, as evidenced by the strong relationship between decreases in nutrient loads and concomitant increases in *Ne* (86,87) (Supplementary Fig. 12, Supplementary Fig. 13; see “Supplementary Discussion” in Supplementary Materials).

In conclusion, we found that two populations of an economically important fish species have had large, simultaneous increases in effective population sizes both historically and over contemporary time scales. The two populations are separated by over 350 km and are genetically divergent from one another, such that they represent two distinct evolutionary lineages (Fig. 1b).

We also found evidence of parallel adaptation to reduced visual clarity in each of the habitats, which may also have been further magnified by centuries-long anthropogenic disturbances and pollution, though we suspect that this genetic adaptation occurred over a longer time scale. The contemporary estimates of *Ne* are orders of magnitude lower than historic estimates, suggesting that recent anthropogenic disturbances may have reduced genetic diversity in Green Bay and Muskegon Lake. While the recent increases in *Ne* suggest that the largest threats to these populations may have passed, the overall point estimates remain low. We suggest that conservation and management efforts targeting yellow perch in these regions should strive to maintain genetic diversity and obtain accurate estimates of population abundances through time. Taken together, these results suggest that the synchrony of key ecological and evolutionary processes can drive parallel demographic and evolutionary trajectories across independent populations. How frequently such synchrony occurs remains an open question in the fields of ecology and evolution.

## Data availability

All raw data generated for this project are stored in the NCBI Short Read Archive (SRA) PRJNA1037191 Great Lakes yellow perch whole genome sequencing. Additionally, unfiltered and filtered VCF files (as described in Supplementary Table 2) are also available (https://doi.org/10.6084/m9.figshare.24517471.v1). The scripts used in this project are hosted in a public repository (https://doi.org/10.6084/m9.figshare.27830115.v1).

## Acknowledgements

We thank the Michigan Department of Natural Resources, the Indiana Department of Natural Resources, the Wisconsin Department of Natural Resources, the Illinois Natural History Survey, and the Grand Traverse Band of Ottawa and Chippewa Indians for assistance with sample collection. In particular, we thank Charles Roswell, Troy Zorn, Tammie Paoli, Dave Clapp, Pat O’Neil, Dave Fielder, David Schindelholz, Erik Olsen, Jayson Beugly, Josey Cline, Taylor Senegal, Greg Chorak, and Scott Koenigbauer. Field collections of yellow perch were authorized by issuance of state-specific collector’s permits and approved by Purdue University’s Animal Care and Use Committee protocol 1112000400. We thank Kurt Thompson for assistance with the map of Muskegon Lake. Secchi depth data were collected and provided by Sarah Bartlett and the NEW Water monitoring program in Green Bay and Sarah Stein and Les Warren of Purdue University. This work was funded by the Great Lakes Fishery Commission (Project ID: 2018_CHR_44072) with additional support from the Great Lakes Fishery Trust (#1456 and #1952) to sample Muskegon Lake.

## Author contributions

MRC, XY, and TOH designed the project, CES and CRR collected samples with agency assistance, XY performed molecular work, XY, MMS, PTE, TJH, CRR, and MRC analyzed the data, and XY and MRC wrote the manuscript with input from all authors. All authors read and approved the final manuscript.

## Additional information

Supplementary materials are available at XXXXXX.

