## Supplementary materials for "Parallel genetic adaptation amidst a background of changing effective population sizes in divergent yellow perch (*Perca flavescens*) populations"

Proceedings of the Royal Society B: Biological Sciences

Article DOI: 10.1098/rspb

**Supplementary Methods:**

***Yellow perch life history***

Yellow perch have a bipartite life cycle consisting of a dispersive larval stage of 30 to 40 days (1,2), followed by demersal, relatively site-attached juveniles and adult life stages (3,4). Adult yellow perch exhibits relatively high site fidelity to spawning grounds (4–6). Female yellow perch lay their eggs in skeins (7) and, after about 15-20 days of egg development in Lake Michigan, embryos hatch into minuscule larvae, which disperse passively in the currents for approximately 30-40 days during their early life stages and may eventually adopt a strategy of combined active and passive dispersal as larvae develop and perform better at swimming (8). Similar to many marine fishes, yellow perch have high fecundity, high larval and juvenile mortality, and large population sizes (9–11). With a relatively short lifespan of about seven years, yellow perch reach maturity at age 1 or 2 in male and at age 2 or 3 in female (12,13). The best estimates for yellow perch generation time are four years +/- one year (13).

***Whole-genome sequencing***

We first extracted DNA from 210 yellow perch collected from seven sample sites in Lake Michigan (Supplementary Table 1) using QIAGEN DNeasy® Blood & Tissue Kits. We tested DNA degradation and potential contamination by running agarose gel electrophoresis and quantified DNA concentration using Qubit 2.0. Genomic DNA was fragmented to an average of ~350 bp through sonication. DNA fragments were then end-polished, A-tailed, and ligated with full-length adapters of Illumina sequencing. We amplified the ligated products with P5 and indexed P7 oligos through PCR, purified PCR products using AMPure XP system, checked the size distribution of libraries by Agilent 2100 Bioanalyzer (Agilent Technologies, CA, US), and quantified the libraries using real-time PCR. The libraries were sequenced on an Illumina

Novaseq 6000 (Illumina Inc., San Diego, CA, USA) to generate 150 bp paired-end reads. Q20 and Q30 (*i.e.*, the base count of Phred values greater than 20 and 30, respectively, divided by the total base counts) ranged from 94.82%-97.32% and 88%-93.12% of all reads, respectively.

***Variant calling and filtering***

We called variants following the pipeline "Germline short variant discovery (SNPs + Indels)" provided by the Genome Analysis Toolkit 4.1.9.0 (GATK 4.1.9.0)(14). The DNA sequencing (DNA-Seq) reads were first mapped to the yellow perch genome (15) using BWA-MEM, BAM files were converted into SAM files sorted by query name using Picard tools, and duplicates were marked using the GATK tool, MarkDuplicatesSpark. Next, we called variants per sample using the GATK tool, HaplotypeCaller, in which we set -ERC to GVCF, --max-alternate-alleles to 100 and --max-num-haplotypes-in-population to 256. Lastly, the GVCF files generated from HaplotypeCaller across all 210 samples were consolidated by chromosome using the GATK tool, GenomicsDBImport, and joint genotyping was conducted using the GATK tool, GenotypeGVCFs, by setting --max-alternate-alleles to 100. We filtered variants using the GATK tool, VariantFiltration, by setting -window to 35 and -cluster to 3 and applying filters "FS >30.0" and "QD < 2.0", and removed indels to retain only single nucleotide polymorphisms (SNPs).

We filtered SNPs using VCFtools (0.1.16). Study-wide MAF filtering is useful for delineating populations, while within-group filtering is essential for identifying rare variants within each population. To create two sets of SNPs for downstream analyses, we first set genotypes called at loci covered by fewer than three reads in each sample as missing values by setting --minDP to 3, deleted SNPs that were genotyped in fewer than 90% of samples by setting --max-missing-count to 42, kept only biallelic SNPs by setting the flags --max-alleles and --min-alleles to 2, and filtered out SNPs with minor allele frequency (MAF) smaller than 0.05 across 210 samples in all seven sample sites to create the **study-wide** data set. This set of SNPs were used to create phylogenetic trees using RAxML version 8.2.12 (16), run principal coordinate analysis using *hierfstat* in R version 4.2.1 (17) and GenAlEx 6.51b2 (18), and conduct population structure analysis on all sample groups using Admixture 1.3 (19) (Supplementary Table 2). Second, we split the VCF file with SNPs by sample site into seven separate VCF files, one for each sample site, set genotypes called at loci covered by fewer than three reads in each sample as missing values by setting --minDP to 3, deleted SNPs that were genotyped in fewer than 90% of samples by setting --max-missing-count to 5 (for Michigan City) or 6 (for the other six sample sites) due to different sample sizes of seven sample sites (Supplementary Table 1), kept only biallelic SNPs by setting the flags --max-alleles and --min-alleles to 2, and filtered SNPs by MAF for each population by discarding SNPs that have MAF smaller than 0.02 or 0.05 within each population to create the **within-group** data sets. The set of SNPs filtered by MAF ≥ 0.02 were used to calculate Tajima's D using VCFtools (0.1.16), while the set of SNPs filtered by MAF ≥ 0.05 were used to assess minor allele frequency distributions, calculate genetic diversity (*i.e.*, observed heterozygosity), estimate effective population size using NeEstimator V2 (20), and reconstruct demographic histories using GONE (21) (Supplementary Table 2). To calculate pairwise $F_{ST}$ for pairs of populations (*i.e.*, Green Bay or Muskegon Lake *vs.* five main basin populations), we created a VCF file containing samples and their genotypic information in the two corresponding populations and filtered the VCF file following the same filtering processes for the within-group data set with MAF ≥ 0.05 across all samples in the two populations in comparison.

### *Population structure analysis*

To infer the phylogenetic relationship among the seven sample sites, we constructed phylogenetic trees using 5,000 randomly selected SNPs. We first converted SNPs in VCF format into PHYLIP format and then constructed a phylogenetic tree using RAxML version 8.2.12 (16). When running RAxML, we selected the GTRGAMMA model, set the random number seed for parsimony inferences to 12345 and the number of alternative runs on distinct starting trees to 1000, turned on rapid bootstrapping, ran rapid bootstrap analysis, and searched for the best-scoring ML tree in each run.

### *Effective population size and demographic history*

In estimating effective population size with 30,000 SNPs randomly selected from the within-group data sets obtained in the current study (Supplementary Table 2; replicated 3 times per population) for each population using the linkage disequilibrium method, as implemented in NeEstimator V2 (20), we selected the random mating model, set the critical value to 0.05, and picked across chromosomes as the locus pairing mode to restrict the analysis to pairs of loci on distinct chromosomes. We further estimated effective population size with 3,000 randomly selected SNPs from a previously obtained RAD-Seq data set (3), again replicated 3 times per sample site for each of 12 sample sites (five from Green Bay, two from Grand Haven, two from Michigan City, one from Milwaukee, one from Naubinway, one from Sutton’s Bay), using the same options described above. The RAD-Seq data set was only used in this particular analysis (Supplementary Table 2) and was employed to validate our findings with additional sample sites and independent extraction, sequencing, and genotyping pipelines.

***Candidate genes and gene ontology (GO) hierarchy networks underlying rapid adaptation***

We created ten separate $F_{ST}$ files: five for each comparison between Green Bay and each of the five main basin sample sites (*i.e.*, Grand Haven, Michigan City, Milwaukee, Naubinway, Sutton's Bay) and five for each comparison between Muskegon Lake and each of the five main basin sample sites. In all $F_{ST}$ output files separately, we first identified SNPs that had $F_{ST} \geq 0.6$. We next took 10 KB steps in either direction of that SNP to identify all SNPs in that region. If additional SNPs in that region continued to have $F_{ST} \geq 0.6$, we continued to expand the window until no additional SNPs of high $F_{ST}$ were found. For each comparison separately, we required that every window had a minimum of 50 SNPs, at least one of which had to have $F_{ST}$ of 0.6 or higher. This step was implemented to filter out regions of the genome with very few segregating sites, which can indicate alignment, genome-assembly, or other spurious errors. We compared all windows to find windows that overlapped among the ten pairwise comparisons. To retain an outlier window, we required that windows from a minimum of eight out of the ten pairwise population comparisons shared at least one high $F_{ST}$ SNP (note that despite this minimum requirement, all windows had at least 82 high $F_{ST}$ SNPs).

***Water clarity, water quality, and phenotypic measurements***

Data on water clarity from Green Bay were provided for four offshore reference sites by the NEW Water monitoring project (www.newwater.us). Data on water clarity for Muskegon Lake and Lake Michigan were collated from past and ongoing projects in the Höök Laboratory (22). Sample sites in Lake Michigan were located near large river mouths, but Secchi readings were taken outside of the river mouth plume. Therefore, these values represent conservatively low estimates of water clarity compared to offshore regions of Lake Michigan. In all cases, Secchi depth was collected using best practices of dropping a black and white Secchi disk on the shaded side of the boat and recording the depth in meters that it is no longer visible.

To collect yellow perch from the littoral zone of Muskegon Lake, boat electrofishing occurred at randomly selected sites around the perimeter of the lake in August 2022. Each site was electrofished for 20 minutes or until a total of 40 yellow perch of at least 150 mm in total length were collected. To collect yellow perch from Lake Michigan, the Michigan Department of Natural Resources set 4×2000 m experimental stretch-mesh gill nets (3.81-15.24 cm) in 9-50 m of water overnight for 12.5-15 hours near the port of Grand Haven during May 2023. To collect yellow perch from Green Bay, the Michigan Department of Natural Resources set 4×97.5 m experimental stretch-mesh gill nets (2.54-12.7 cm) in 6.4-9.5 m of water overnight for approximately 24 hours near the Menominee River during September 2023. Yellow perch from both locations were transported to the laboratory where total body length was measured. Fish were individually marked (with a paper tag), fixed in 10% formalin for six days, and then transferred to 70% ethanol for long-term preservation. After 60 days in ethanol, eye diameter was measured using electronic calipers. To compare eye diameter among the three locations, we analyzed yellow perch with total lengths of 150-250 mm to ensure comparisons of yellow perch with similar sizes ($n=31$ for Menominee in Green Bay, $n=39$ for Muskegon Lake, and $n=27$ for Grand Haven in Lake Michigan). We only collected fin clips from the 210 fish we sequenced, thus the fish measured for eye size were not the same set of fish that were genotyped.

**Supplementary Results:**

***Genetic divergence among yellow perch sample sites***

The high genetic divergence between Green Bay and main basin sample sites, as revealed by Miller (23) and Schraidt *et al.* (3), could be driven by the limited larval exchange between the two basins (24). One sample collected from Muskegon Lake (sample id: D9332) in July 2016 clustered with main basin sample sites, suggesting that migration between the main basin and Muskegon Lake is possible. Interestingly, D9332 was captured via a nighttime boat electrofishing survey at the State Park site in Muskegon Lake (see Fig. 1 in Bhagat and Ruetz 2011 (25)), which is closest to Lake Michigan among the four sites we sampled in Muskegon Lake. Additionally, D9332 was 10.4 cm TL, which was smaller than the mean size of yellow perch captured in Muskegon Lake (mean = 15.5 cm TL). Therefore, D9332 was likely born in the main basin of Lake Michigan and subsequently moved into Muskegon Lake. The movement of yellow perch from the main basin into Muskegon Lake has been suggested previously by Chorak *et al.* (26,27). Despite the clear movement of a single individual, we found the remaining Muskegon Lake samples to be genetically divergent from main basin sample sites (Fig. 1b,d; see Supplementary Table 5 for all pairwise $F_{ST}$ estimates).

**Supplementary Discussion:**

We found a strong relationship between decreases in nutrient loads and concomitant increases in *Ne* in two geographically separated yellow perch populations (Supplementary Fig. 12, Supplementary Fig. 13). While we are aware that correlation does not equal causation, we speculate that this result may suggest that improvements in water quality can result in increased *Ne*. Water quality in Green Bay and Muskegon Lake was at its worst in the early 1970s, after which environmental remediation resulted in substantial water quality and habitat improvements, particularly for reductions in phosphorus (28,29) (Supplementary Fig. 12). Given an estimated generation time of four years (9,12), effective population sizes of yellow perch populations began rebounding in the early 2000s, some 20-30 years after cleanup efforts were well under way (generation time estimates of three and five years show similar results; Supplementary Fig. 12; Supplementary Fig. 13). For both Green Bay and Muskegon Lake, there is a strong relationship between total phosphorous and effective population size (Supplementary Fig. 12), suggesting that, as nutrient loading decreased, *Ne* increased (but see caveats below). Note that while more data on phosphorus were available for Muskegon Lake due to the use of sediment cores to quantify changes through time, concentrations of phosphorus and other nutrients also have declined in Green Bay since the 1970s (28).

Mismatches may exist between *Ne* estimated by GONE and actual abundances obtained from fishery dependent or independent surveys in some populations. Historical data on catch rate per unit effort (CPUE) suggests that abundance of yellow perch in Green Bay started increasing since late 1970s and peaked around 1980s (30), while *Ne* of the Green Bay site we sampled remained relatively low through 1970s and 1980s, started increasing around 1990s, and reached maximum around 2000 (Supplementary Fig. 12, Supplementary Fig. 13). There are several potential explanations for this mismatch. First, the timing of major historical events estimated by GONE may not be accurate. Although GONE is fairly robust to genotyping errors, temporal heterogeneity of samples, population admixture, and population subdivision, biases arising from violations of isolation and panmixia of populations under study may not be fully eliminated (21,31). Second, overlapping generations and fishing-induced evolution may both complicate the estimation of generation times. Third, slight differences in *Ne* are detected across sample sites in Green Bay (Fig. 3b), so the reconstructed demographic history for our collection site may not be representative of *Ne* of the entire region and thus may not be representative for all of Green Bay. Lastly, current methods for estimating fish abundance are also not free of bias. CPUE may not be a reliable estimator for stock abundance because: 1.) the correlation between commercial catch rates and stock abundance may not hold due to aggregated fishing, undocumented changes in fishing efforts, differential proportions of species captured, and by-catch discard (32–35), and 2.) insufficient time series of data, owing to the difficulty in data collection over a large temporal scale, may reduce confidence in estimates of abundance derived from statistical models (36).

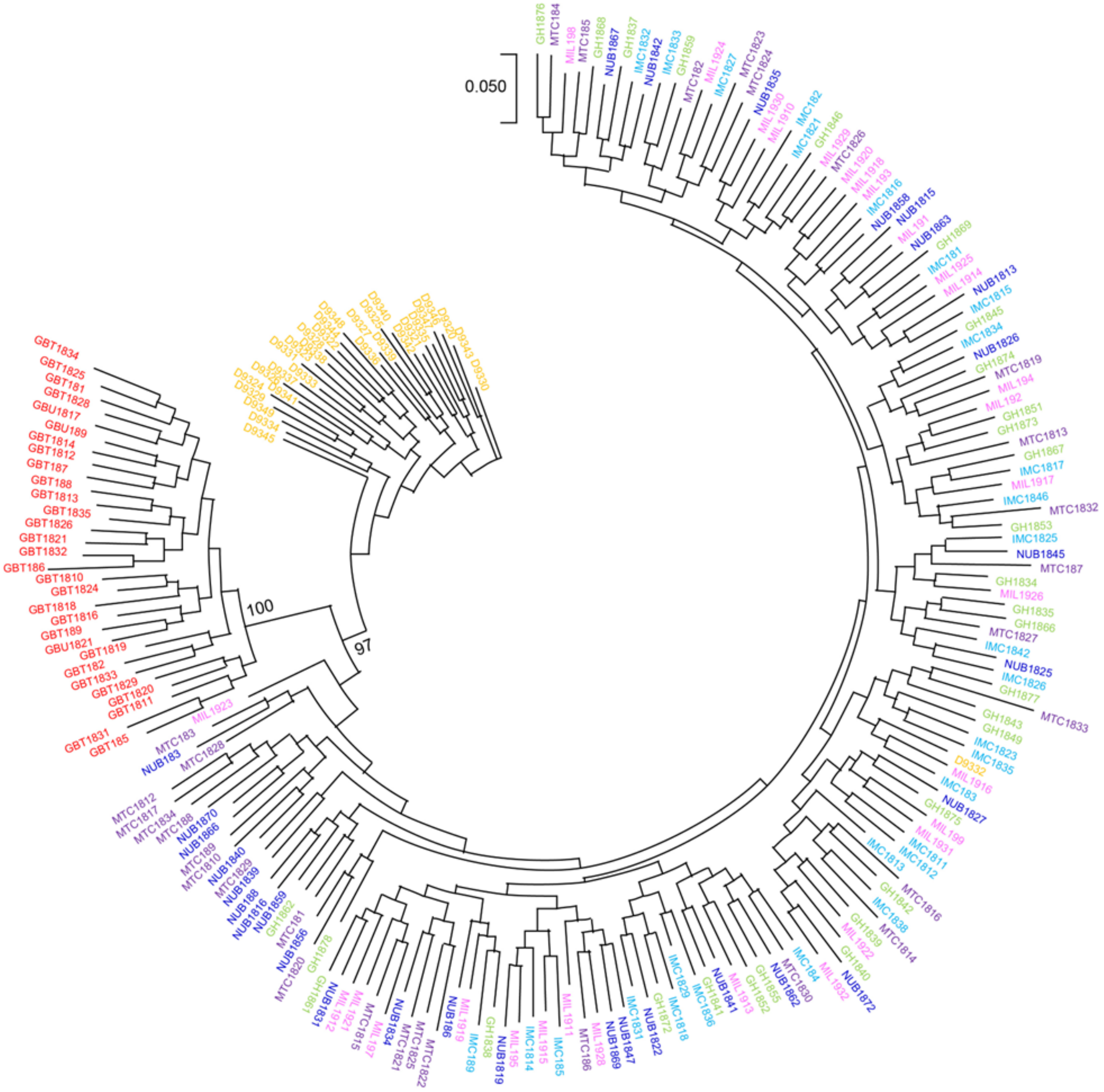


● Green Bay ● Muskegon Lake ● Grand Haven ● Michigan City ● Milwaukee ● Naubinway ● Sutton's Bay

**Supplementary Figure 1.** Phylogenetic tree for all 210 samples. The phylogenetic tree was constructed with 5,000 randomly selected SNPs from all 210 samples using RAxML version 8.2.12 (16). Numbers on branches indicate the percentage of bootstraps returning the best-scoring ML tree in 1,000 alternative runs with distinct starting trees and values are only shown when greater than 90%.

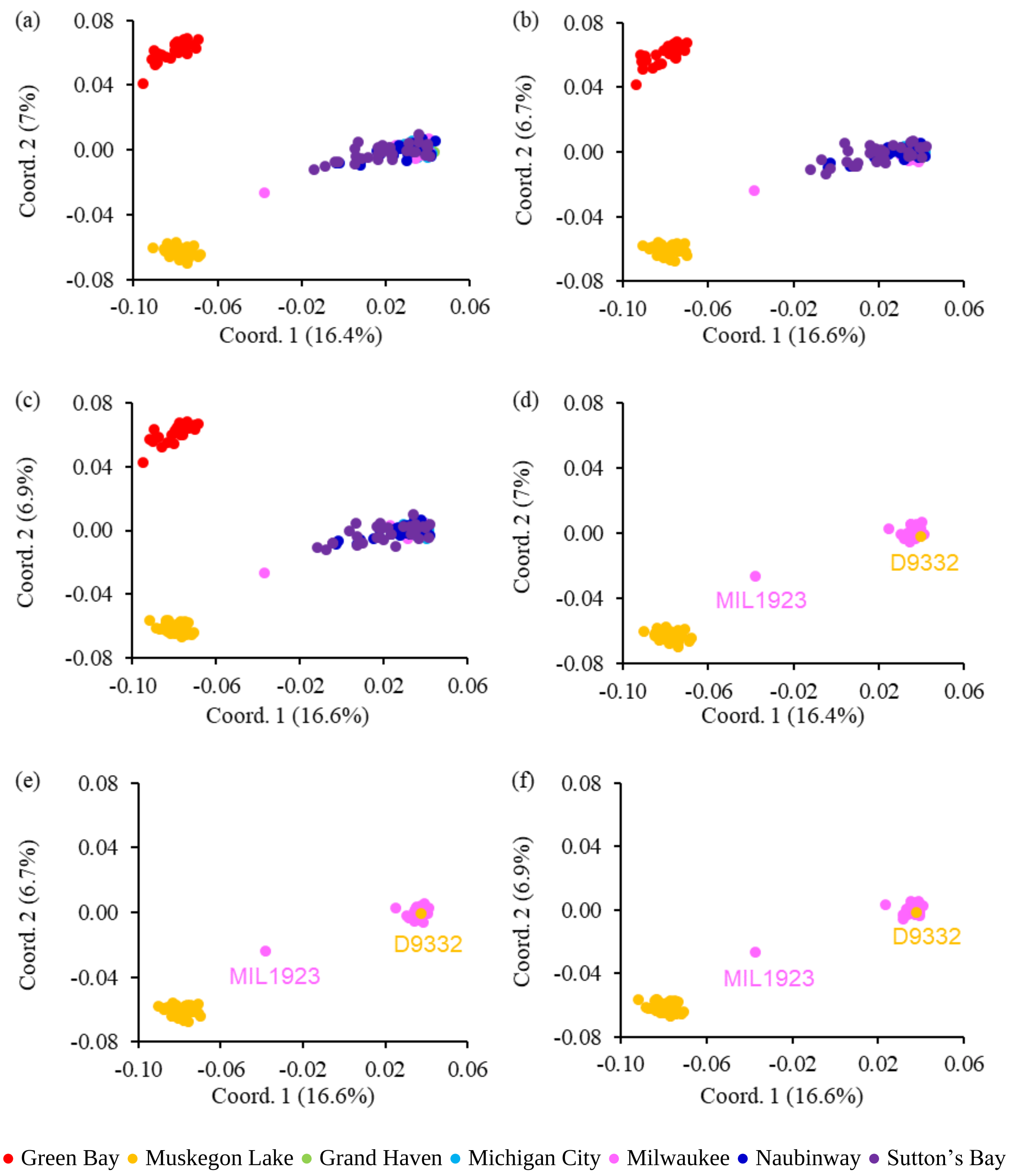


**Supplementary Figure 2.** Principal coordinate analysis using *hierfstat* in R version 4.2.1 for all seven sample sites (a, b, c) and Muskegon Lake and Milwaukee (d, e, f). Principal coordinate analysis was conducted independently three times with three different sets of 50,000 randomly selected SNPs to account for the potential variation caused by different SNPs. One Muskegon Lake sample clusters with the main basin populations, suggesting potential migration from the main basin. One Milwaukee sample falls comparatively further away from the main basin populations, suggesting potential gene flow or shared ancestry with a fish of unknown origin. Panel (a) is the same as Fig. 1c in the main text.

(a)

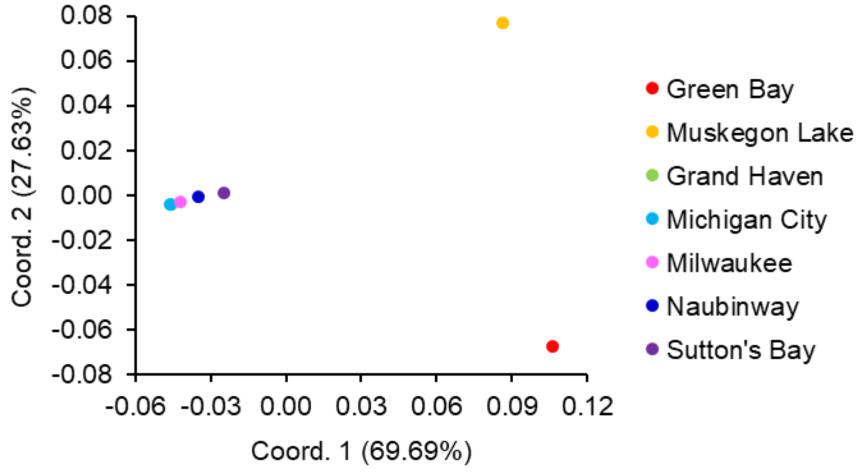


(b)

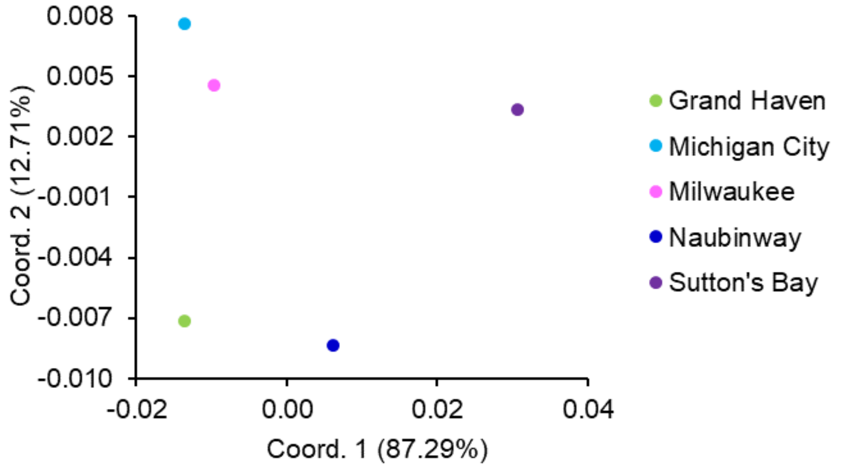


**Supplementary Figure 3.** Principal coordinate analysis of mean pairwise $F_{ST}$ for all seven sample sites (a) and five main basin sample sites (b) based on mean Weir and Cockerham's $F_{ST}$ between pairs of sample sites (see Supplementary Table 5 for details). Note that the northern main basin sample sites are slightly divergent from the southern main basin sample sites (axis 1 of panel b), a finding congruent with earlier RAD-Seq analyses (3), and Grand Haven is missing in panel (a) because it is hidden behind Michigan City.

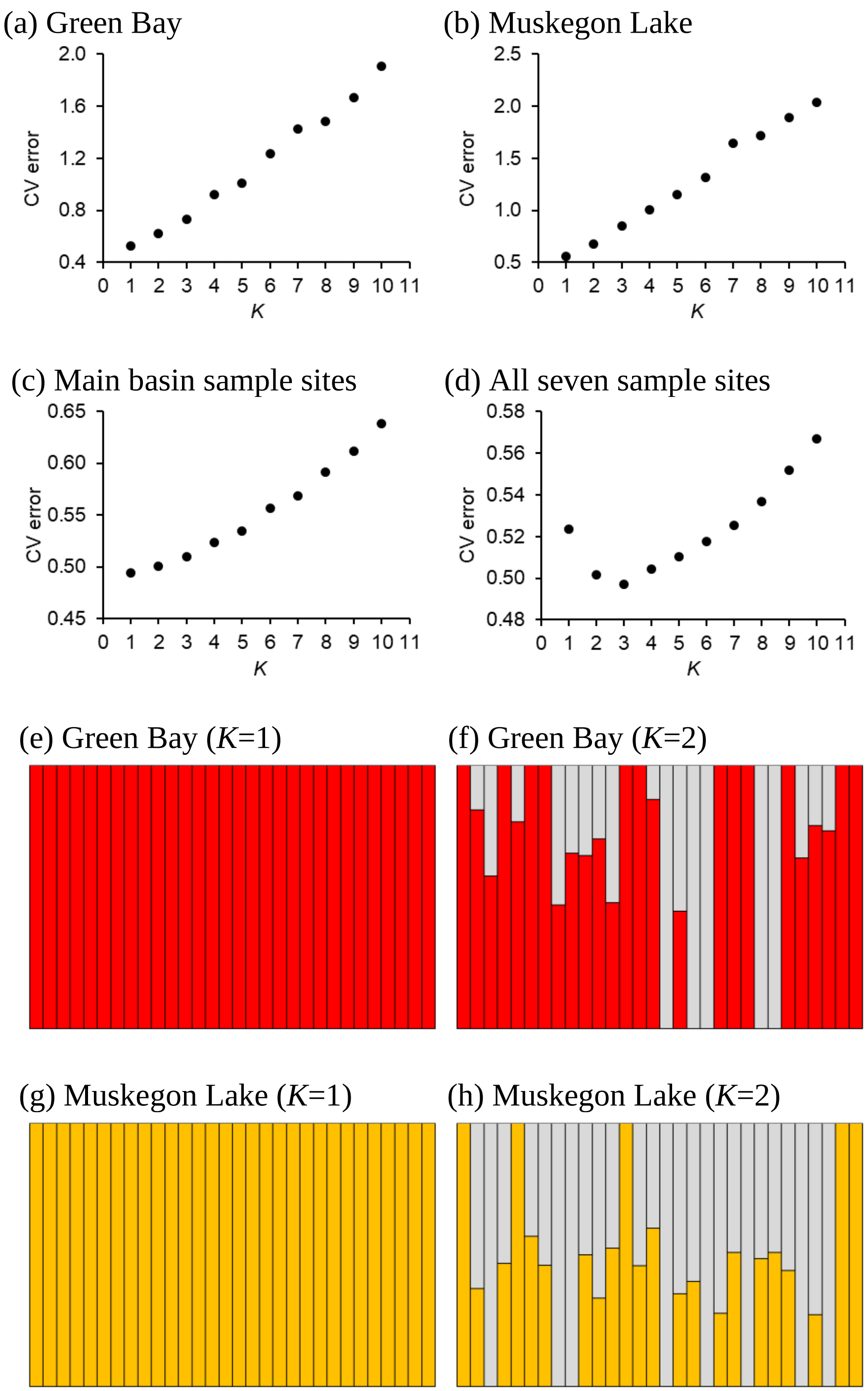


**Supplementary Figure 4.** Optimal *K* in admixture analysis for Green Bay (a, optimal *K*=1), Muskegon Lake (b, optimal *K*=1), main basin sample sites (c, optimal *K*=1), and all seven sample sites (d, optimal *K*=3) and ancestry plots of Green Bay (e, f) and Muskegon Lake (g, h) populations.

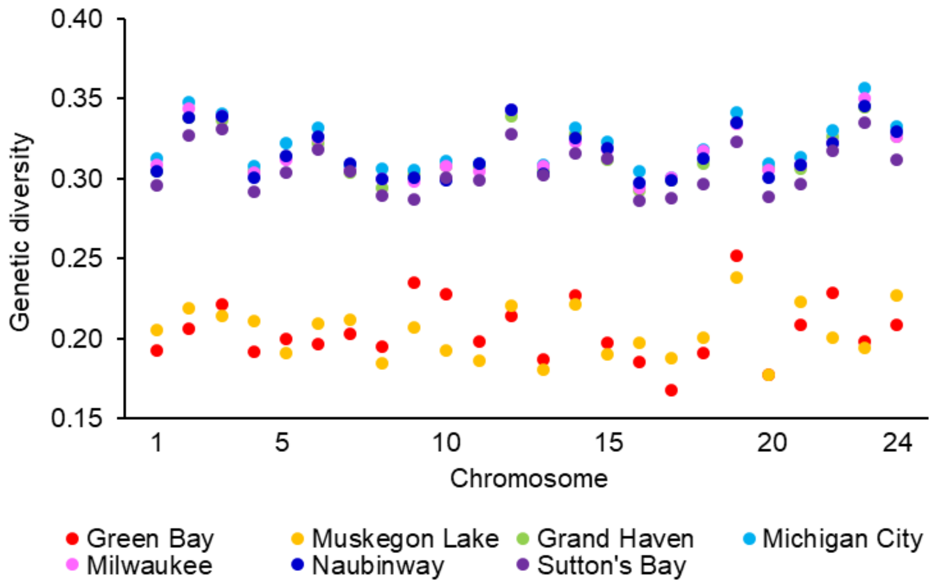


**Supplementary Figure 5.** Chromosome-level genetic diversity as measured by observed heterozygosity. Observed heterozygosity was calculated as the number of heterozygous individuals divided by the number of genotyped individuals at SNPs from the within-group data set (MAF ≥ 0.05) and averaged across all SNPs within each chromosome.

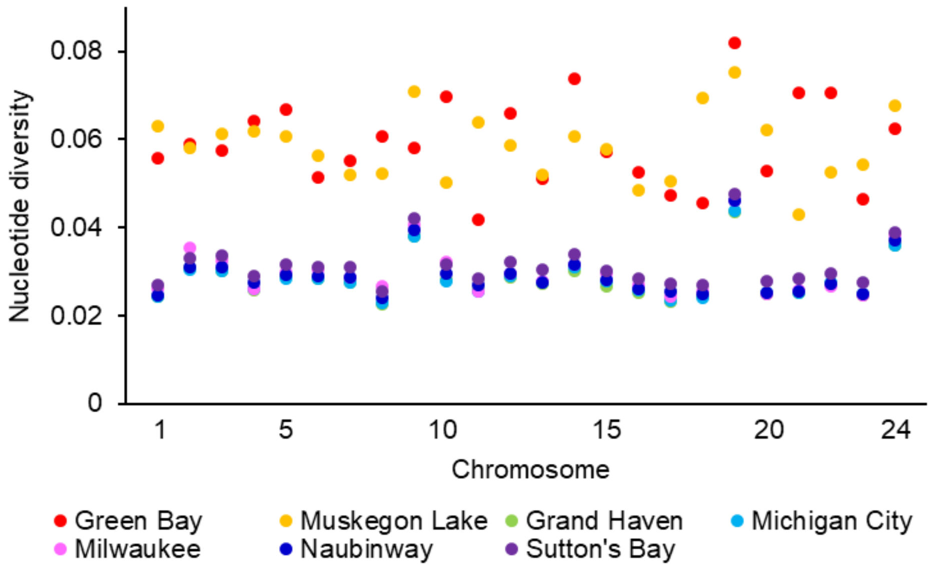


**Supplementary Figure 6.** Nucleotide diversity (π) of Green Bay, Muskegon Lake, and Lake Michigan main basin populations calculated with biallelic sites that are covered by at least three reads in each sample (including both polymorphic and invariant sites) using PIXY (37).

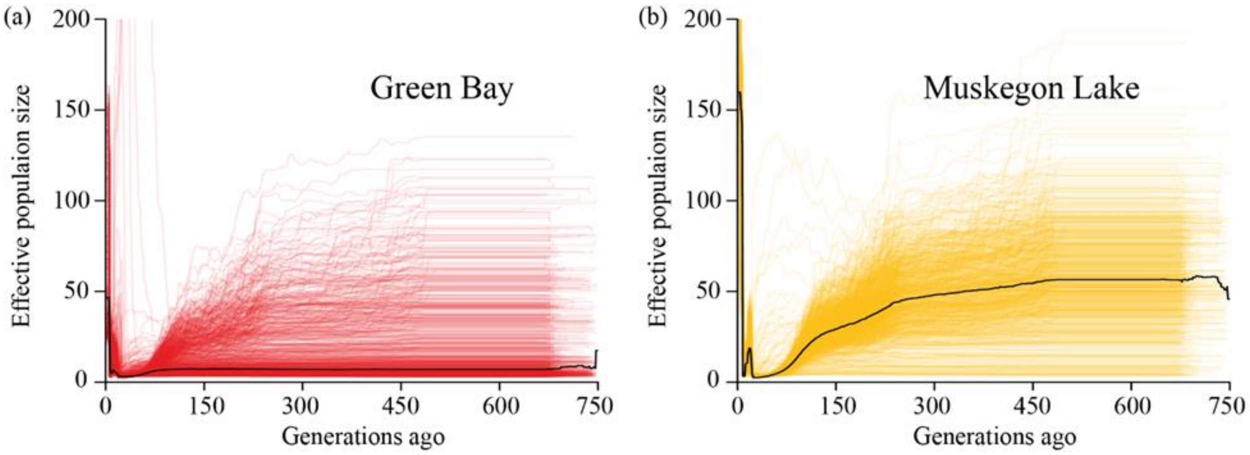


**Supplementary Figure 7.** Demographic histories of Green Bay (a, red) and Muskegon Lake (b, orange) reconstructed with 1,000 replicates of 30,000 SNPs randomly selected from the within-group data set using GONE with a black line indicating median effective population size. In comparison to Fig. 3d,e in the main text, (a) and (b) illustrate the entire demographic inferences of Green Bay and Muskegon Lake over the past 750 generations, respectively. Note, however, that GONE is not very accurate in reconstructing demographic histories prior to bottlenecks (38).

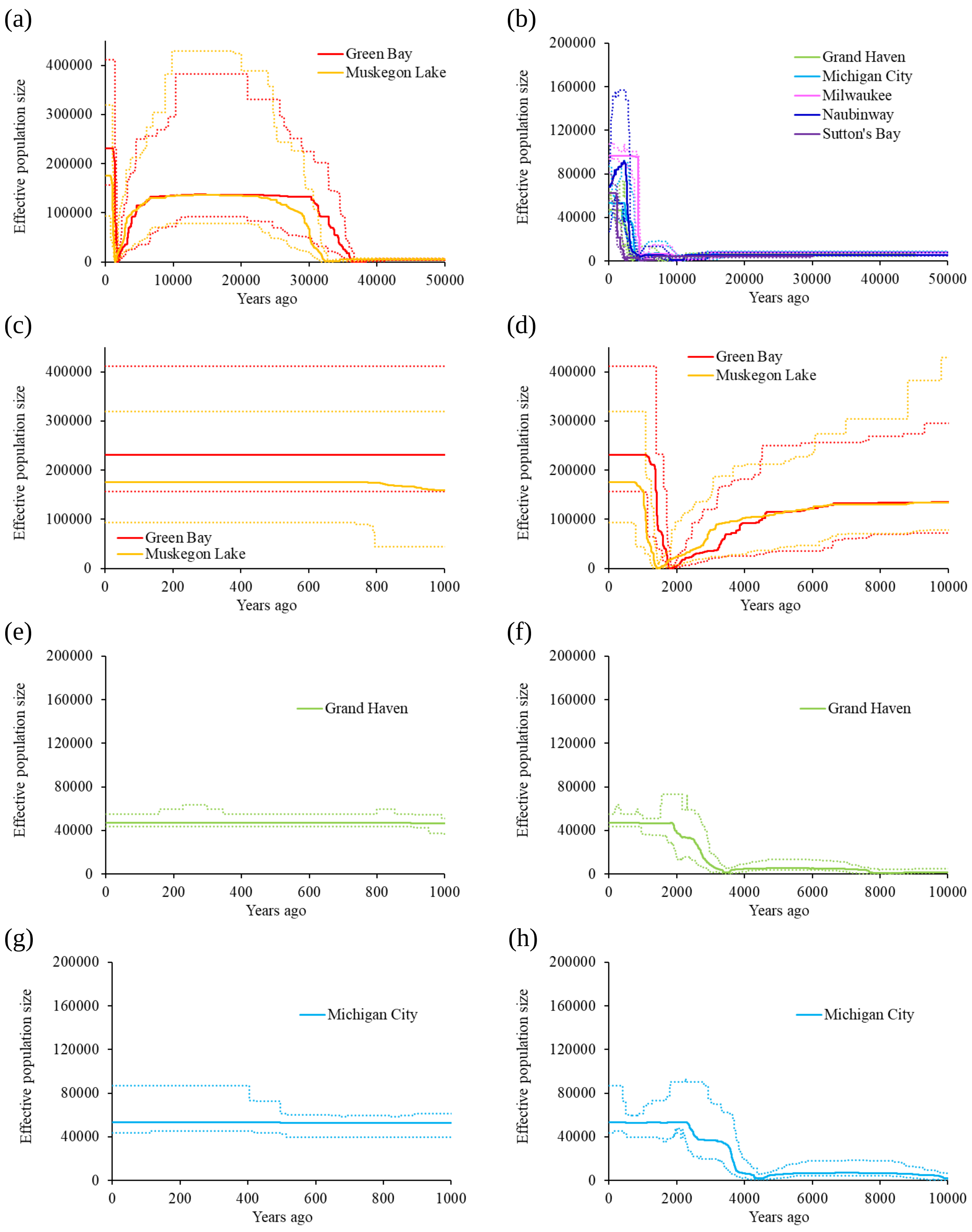

(a)
Effective population size
Years ago
Green Bay
Muskegon Lake
(b)
Grand Haven
Michigan City
Milwaukee
Naubinway
Sutton's Bay
(c)
(d)
(e)
(f)
(g)
(h)

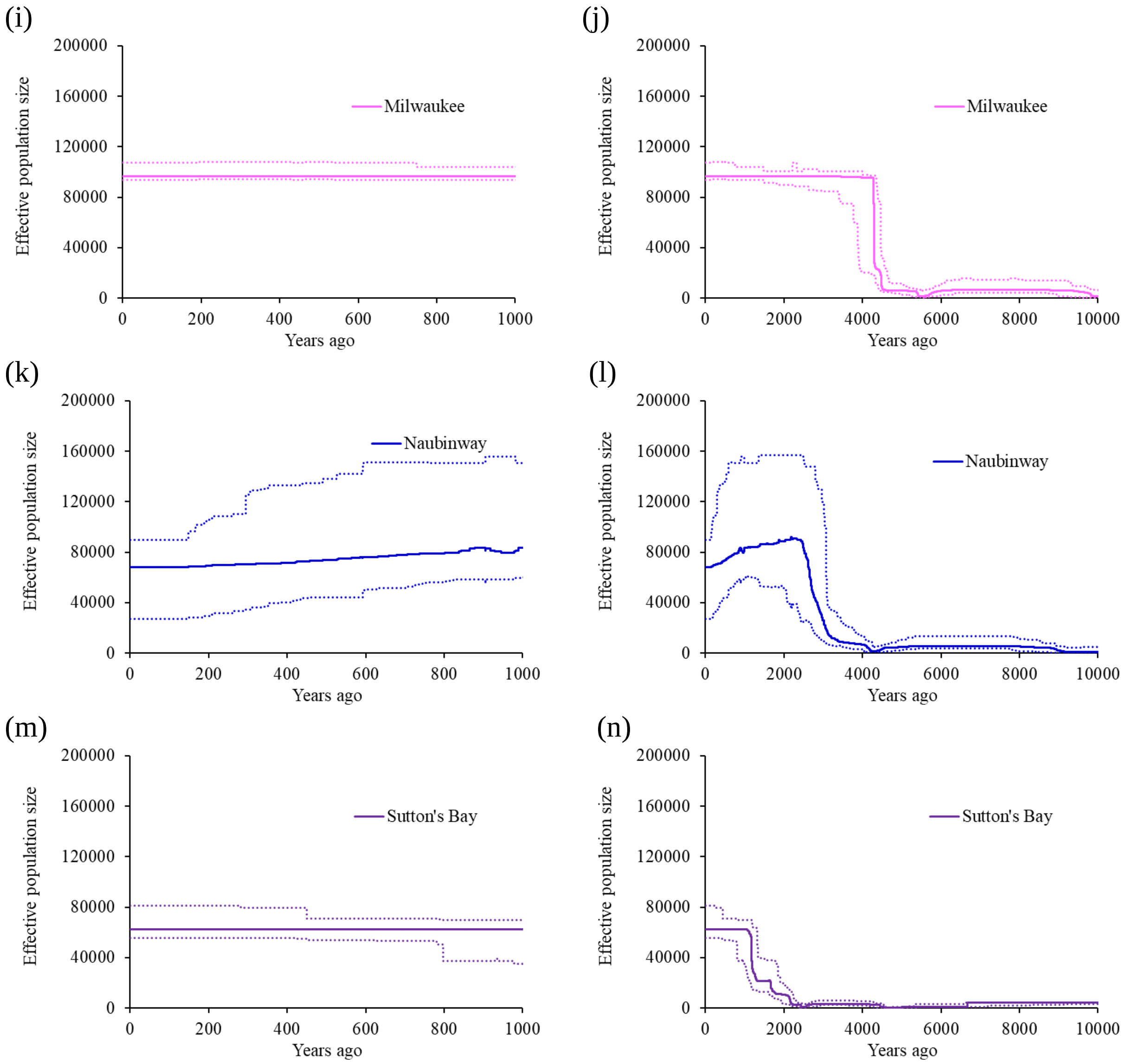


**Supplementary Figure 8.** Demographic histories reconstructed by Stairway Plot 2 (39) with a generation time of three years with solid lines indicating median *Ne* and dashed lines indicating 95% CI of *Ne*.

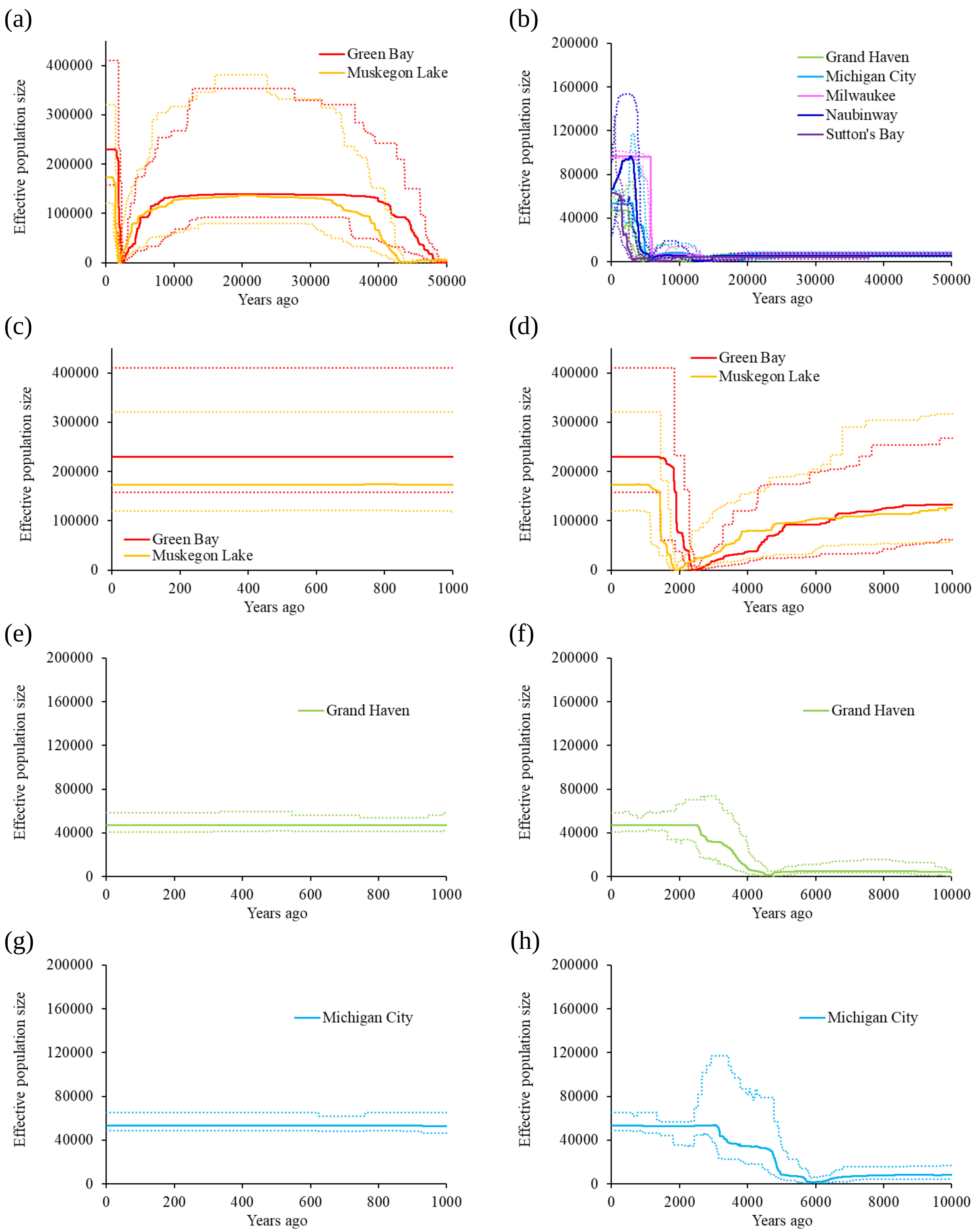

(a)
Effective population size
Years ago
Green Bay
Muskegon Lake
(b)
Grand Haven
Michigan City
Milwaukee
Naubinway
Sutton's Bay
(c)
(d)
(e)
(f)
(g)
(h)

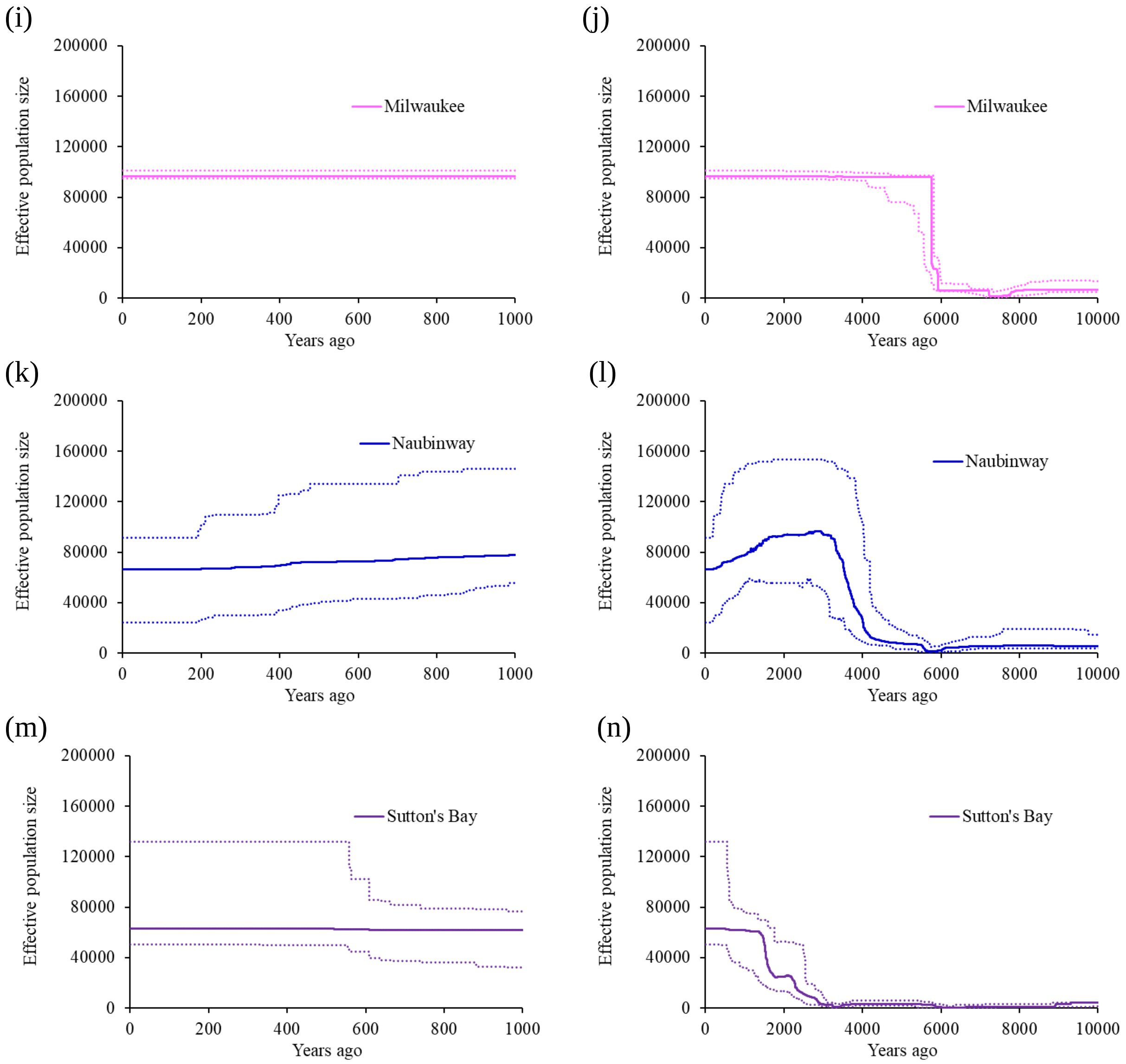


**Supplementary Figure 9.** Demographic histories reconstructed by Stairway Plot 2 (39) with a generation time of four years with solid lines indicating median *Ne* and dashed lines indicating 95% CI of *Ne*.

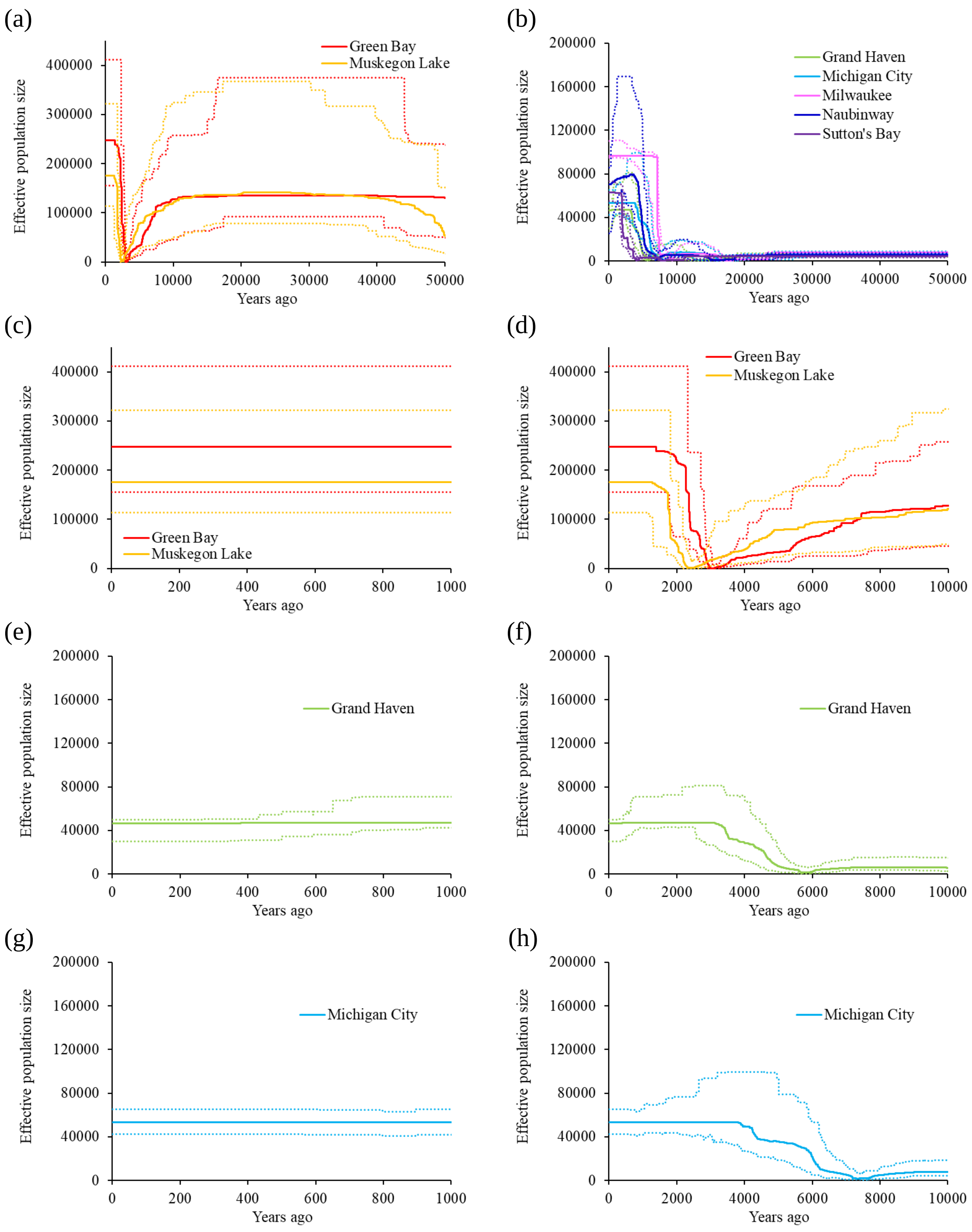
(a)
Green Bay
Muskegon Lake
Effective population size
Years ago
(b)
Grand Haven
Michigan City
Milwaukee
Naubinway
Sutton's Bay
Effective population size
Years ago
(c)
Green Bay
Muskegon Lake
Effective population size
Years ago
(d)
Green Bay
Muskegon Lake
Effective population size
Years ago
(e)
Grand Haven
Effective population size
Years ago
(f)
Grand Haven
Effective population size
Years ago
(g)
Michigan City
Effective population size
Years ago
(h)
Michigan City
Effective population size
Years ago

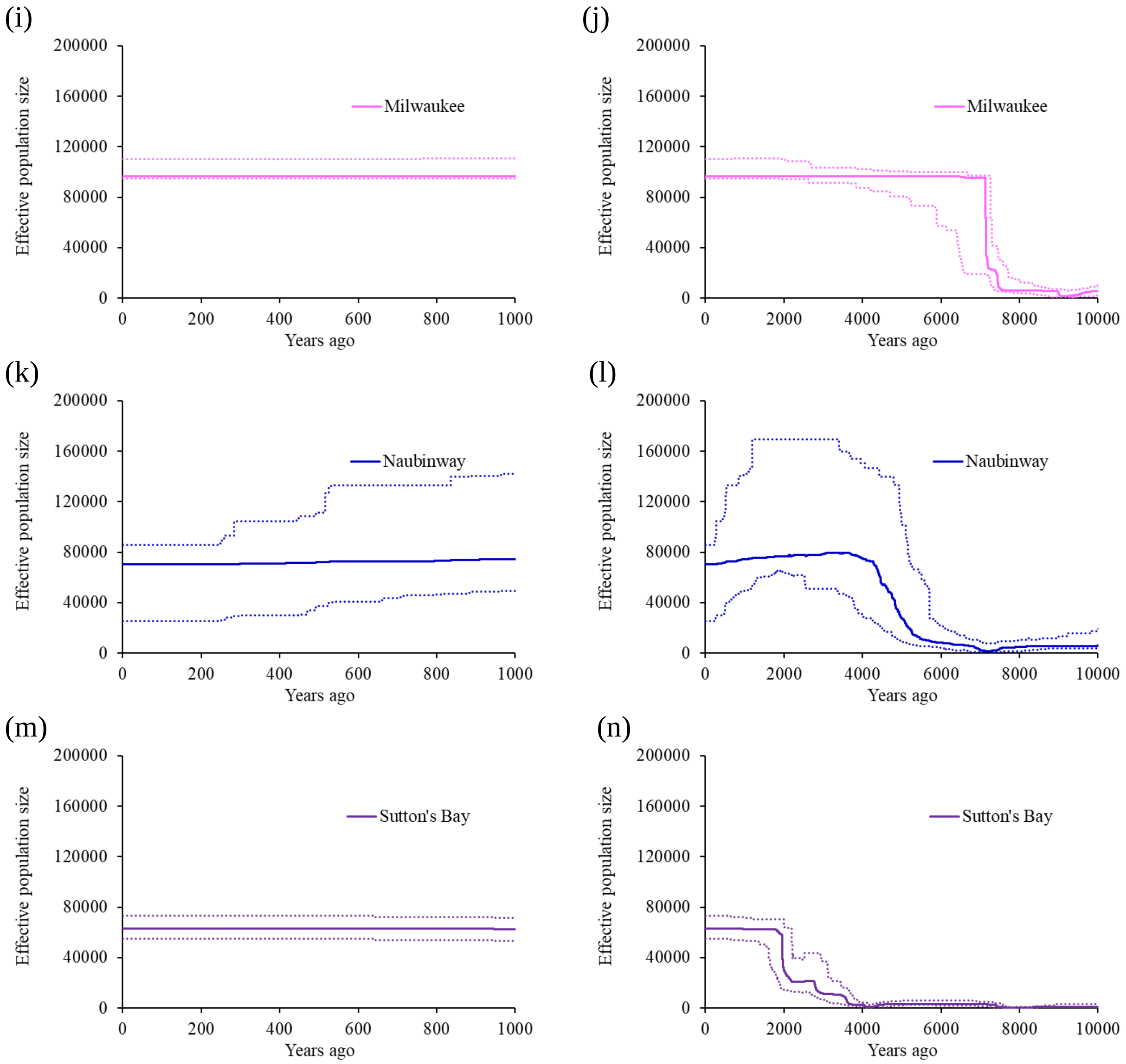


**Supplementary Figure 10.** Demographic histories reconstructed by Stairway Plot 2 (39) with a generation time of five years with solid lines indicating median *Ne* and dashed lines indicating 95% CI of *Ne*.

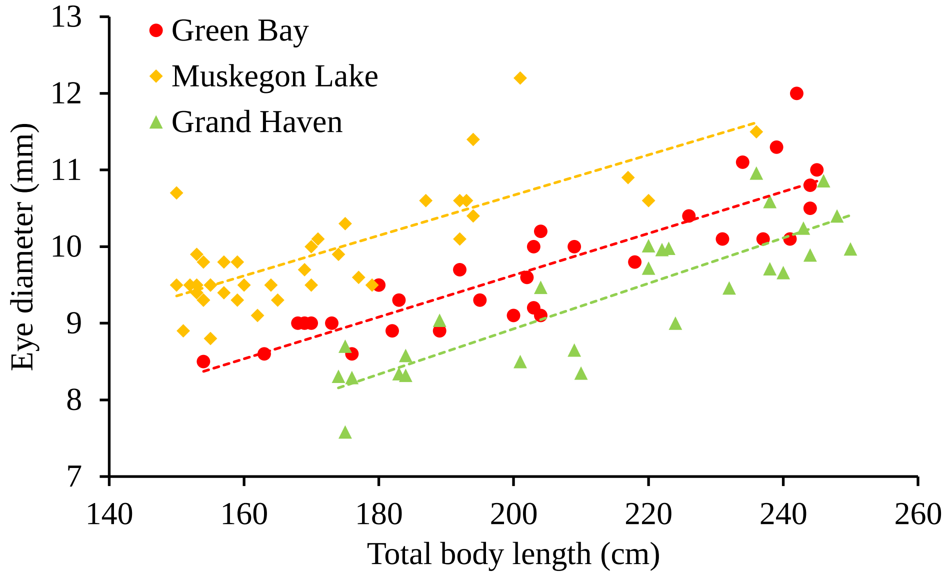


**Supplementary Figure 11.** Regression of eye diameter (mm) on total body length (cm) of yellow perch collected from Menominee in Green Bay, Muskegon Lake, and Grand Haven (*i.e.*, Lake Michigan main basin). A significant positive relationship is detected between eye diameter and total body length ($P < 0.001$), but slopes of linear regressions of eye diameter on total body length are not significantly different for yellow perch among Green Bay, Muskegon Lake, and Grand Haven ($P = 0.779$).

(a) (b)

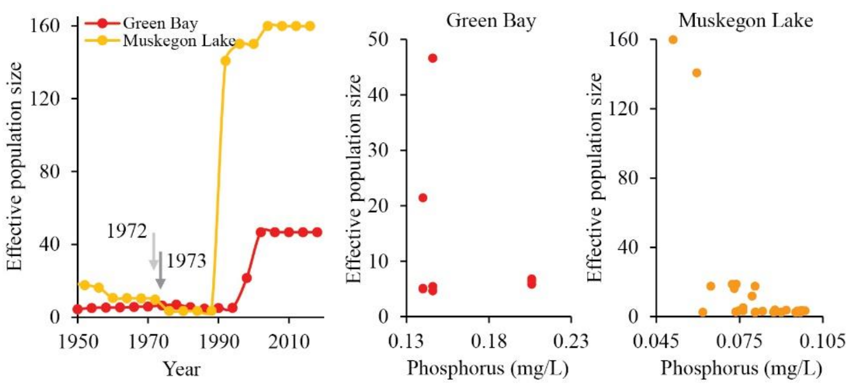


**Supplementary Figure 12.** Increases in effective population size are correlated with environmental remediation. Along with the ecosystem restoration driven by environmental initiatives, such as the Clean Water Act since 1972 (indicated with a light gray arrow) and the Muskegon County advanced wastewater treatment since 1973 (indicated with a dark gray arrow), effective population size started rebounding shortly after policies were implemented (a; generation time=four years). Increases in effective population sizes are associated with substantial reductions in total phosphorus in both Green Bay and Muskegon Lake (b; generation time=four years). Data on total phosphorus were taken from Fig. 2 (*i.e.*, average TP) in Harris et al. (28) for Green Bay and were extracted from Fig. 5c in Liu et al. (29) using WebPlotDigitizer (https://automeris.io/WebPlotDigitizer) (40) for Muskegon Lake.

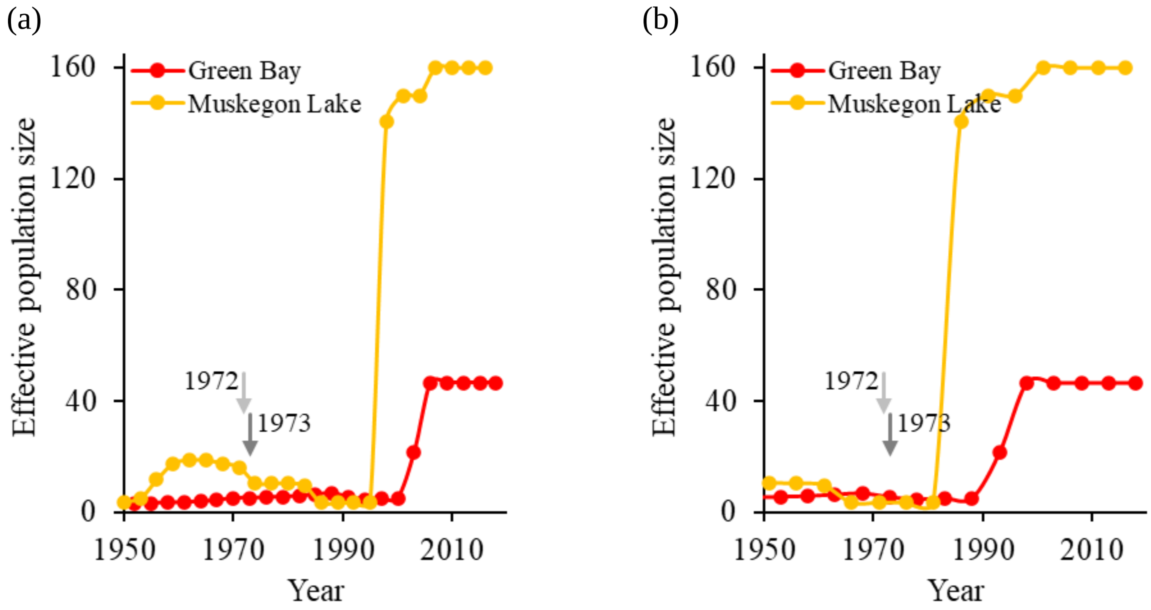


**Supplementary Figure 13.** Effective population sizes of Green Bay and Muskegon Lake before and after environmental remediation driven by environmental initiatives, such as the Clean Water Act since 1972 (indicated with a light gray arrow) and the Muskegon County advanced wastewater treatment since 1973 (indicated with a dark gray arrow). Given an estimated generation time of three (a) or five (b) years, the effective population sizes of yellow perch populations began rebounding approximately 10-30 years after clean-up efforts were under way.

**Supplementary Table 1.** Sampling information where we report the sample site name, the year and month in which sampling was conducted (sampling month is available for a subset of samples), the number of individuals sequenced, and the latitude and longitude (degree).

| Sample site | Sample date | Sample size | Latitude | Longitude |
|---|---|---|---|---|
| Green Bay | 2018 (May) | 30 | 44.513332 | -88.015831 |
| Muskegon Lake | 2016 (July) | 30 | 43.224194 | -86.235809 |
| Grand Haven | 2018 | 30 | 43.063073 | -86.228386 |
| Michigan City | 2018 (May, June) | 29 | 41.707539 | -86.895030 |
| Milwaukee | 2018 (November), 2019 (May, June) | 30 | 43.038902 | -87.906471 |
| Naubinway | 2018 | 30 | 46.092497 | -85.447597 |
| Sutton’s Bay | 2018 (June) | 31 | 44.976666 | -85.650639 |

**Supplementary Table 2.** Differently filtered data sets of SNPs for population genetic and demographic analyses where we report the specific analysis for which different data sets were used and how the corresponding results are illustrated. We filtered SNPs by minor allele frequency (MAF) in two ways to create two data sets of SNPs suitable for different analyses, a study-wide data set and a within-group data set. Briefly, we filtered out SNPs with minor allele frequency (MAF) smaller than 0.05 across 210 samples in all seven sample sites to create the **study-wide** data set and filtered SNPs by MAF for each population by discarding SNPs that have MAF smaller than 0.02 (for Tajima's D calculation only) or 0.05 within each population to create the **within-group** data sets. Study-wide MAF filtering is useful for delineating populations, while within-group filtering is essential for identifying rare variants within each population. The RAD-Seq data set from Schraidt *et al.* (3) was used only for additional LD-based estimates of *Ne* and included additional collection sites.

| **Data sets of SNPs** | **Analysis** | **Results** |
|---|---|---|
| Study-wide data set | Phylogenetic tree construction | Fig. 1b<br>Supplementary Fig. 1 |
| Study-wide data set | Principal coordinate analysis | Fig. 1c<br>Supplementary Fig. 2<br>Supplementary Fig. 3 |
| Study-wide data set | Admixture analysis | Fig. 1d<br>Supplementary Fig. 4 |
| Within-group data set | Comparison of numbers of SNPs | Fig. 2a |
| Within-group data set | Comparison of minor allele frequency across sample sites | Fig. 2b |
| Within-group data set (MAF $\geq 0.02$) | Tajima's D calculation | Fig. 2c |
| Within-group data set | Effective population size estimated by NeEstimator V2 | Fig. 3a |
| Within-group data set | Genetic diversity calculation | Fig. 3c<br>Supplementary Fig. 5 |
| Within-group data set | Nucleotide diversity calculation | Supplementary Fig. 6 |
| Within-group data set | Demographic history reconstructed by GONE | Fig. 3d<br>Fig. 3e<br>Supplementary Fig. 7 |
| RAD-Seq data set | Effective population size estimated by NeEstimator V2 | Fig. 3b |

**Supplementary Table 3.** Mapping quality of samples.

| Sample ID | No. total reads | No. mapped reads | No. reads with mapping quality ≥ 20 | No. reads with mapping quality ≥ 60 |
|---|---|---|---|---|
| D9_320 | 68974612 | 68749732 | 62937577 | 57988746 |
| D9_321 | 74566897 | 74351942 | 67920178 | 62324911 |
| D9_322 | 67046756 | 66841947 | 61658345 | 56928066 |
| D9_323_01 | 6641129 | 6624704 | 6095028 | 5614903 |
| D9_323_02 | 65871425 | 65693206 | 60334194 | 55550578 |
| D9_324_01 | 15211359 | 15168756 | 13877336 | 12725733 |
| D9_324_02 | 55650616 | 55485192 | 50653463 | 46422030 |
| D9_325 | 68281196 | 68073383 | 62133017 | 57036603 |
| D9_326 | 73899079 | 73679379 | 67372931 | 61959692 |
| D9_327 | 80464504 | 80157897 | 71445105 | 64967087 |
| D9_328 | 77365549 | 77134390 | 70435287 | 64877972 |
| D9_329 | 67774495 | 67555970 | 62287865 | 57544889 |
| D9_330_01 | 6566099 | 6544459 | 5990365 | 5510866 |
| D9_330_02 | 65534880 | 65300036 | 59647765 | 54830635 |
| D9_331 | 67886493 | 67646208 | 62130041 | 57335329 |
| D9_332 | 76423556 | 76195516 | 70378444 | 65088422 |
| D9_333 | 72882737 | 72657501 | 66559265 | 61333697 |
| D9_334 | 66618680 | 66432719 | 60884016 | 56013353 |
| D9_335 | 68047077 | 67869904 | 62297028 | 57392000 |
| D9_336 | 83353851 | 83108303 | 76491550 | 70484719 |
| D9_337 | 70522338 | 70315012 | 65100977 | 60249553 |
| D9_338 | 79973085 | 79760831 | 73614776 | 68004053 |
| D9_339 | 99880285 | 99575820 | 91851878 | 84831527 |
| D9_340 | 69603336 | 69398869 | 62949563 | 57517590 |
| D9_341 | 68204357 | 68002230 | 62887494 | 58140145 |
| D9_342 | 72957028 | 72740460 | 66847119 | 61551112 |
| D9_343 | 85120740 | 84828815 | 76867347 | 70333007 |
| D9_344_01 | 63201193 | 63028680 | 58089887 | 53543410 |
| D9_344_02 | 7470224 | 7450340 | 6847199 | 6309619 |
| D9_345 | 73593597 | 73384082 | 67378443 | 62050839 |
| D9_346 | 67112446 | 66934763 | 61664982 | 56889246 |
| D9_347 | 67550362 | 67382640 | 61606572 | 56543407 |
| D9_348 | 80185861 | 79964261 | 73681463 | 68021652 |
| D9_349 | 79121017 | 78922319 | 72621400 | 66994348 |
| GBT18_1 | 68623479 | 68357250 | 61317646 | 55922454 |
| GBT18_2_01 | 63692603 | 63491543 | 57758862 | 52877258 |
| GBT18_2_02 | 7283936 | 7262243 | 6584599 | 6027384 |
| GBT18_5 | 71119000 | 70693177 | 60855292 | 54432686 |
| GBT18_6 | 70139708 | 69853831 | 61378131 | 55464634 |
| GBT18_7 | 73787148 | 73531437 | 64150405 | 57812429 |
| GBT18_8 | 84790813 | 84492446 | 76673301 | 69972318 |

| Sample ID | No. total reads | No. mapped reads | No. reads with mapping quality ≥ 20 | No. reads with mapping quality ≥ 60 |
|---|---|---|---|---|
| GBT18_9_01 | 62327039 | 62138895 | 56549703 | 51789542 |
| GBT18_9_01 | 9853830 | 9822696 | 8920447 | 8168135 |
| GBT18_10_01 | 59318041 | 59115344 | 53839765 | 49337484 |
| GBT18_10_02 | 12195698 | 12154728 | 11039034 | 10115936 |
| GBT18_11 | 71910779 | 71659746 | 64183444 | 58483919 |
| GBT18_12 | 69998187 | 69762886 | 63235807 | 57788229 |
| GBT18_13 | 83743156 | 83378753 | 75970165 | 69584654 |
| GBT18_14 | 69609065 | 69367539 | 63063025 | 57689296 |
| GBT18_16 | 79846107 | 79588194 | 72660450 | 66677010 |
| GBT18_18 | 77609825 | 77323320 | 70124655 | 64082735 |
| GBT18_19 | 67033193 | 66780220 | 60583511 | 55427939 |
| GBT18_20 | 86698500 | 86405263 | 78790592 | 72073647 |
| GBT18_21 | 83699226 | 83434882 | 75964802 | 69479441 |
| GBT18_24 | 87663824 | 87328045 | 79514753 | 72812469 |
| GBT18_25 | 84133021 | 83871561 | 76826878 | 70454442 |
| GBT18_26 | 80628626 | 80385665 | 73803392 | 67753037 |
| GBT18_28 | 82991626 | 82681361 | 75308149 | 68969798 |
| GBT18_29 | 98921913 | 98595327 | 89819103 | 82203138 |
| GBT18_31 | 66323385 | 66082811 | 59922001 | 54709298 |
| GBT18_32 | 81387684 | 81151006 | 73932707 | 67705258 |
| GBT18_33 | 73966413 | 73737824 | 67358284 | 61728889 |
| GBT18_34_01 | 6563900 | 6544615 | 5948063 | 5447558 |
| GBT18_34_02 | 64581427 | 64380496 | 58819997 | 53850821 |
| GBT18_35 | 72262119 | 72027236 | 66066040 | 60637058 |
| GBU18_9 | 79845989 | 79592301 | 72729220 | 66601414 |
| GBU18_17 | 84293767 | 83926951 | 76552734 | 70199378 |
| GBU18_21 | 77260952 | 77033508 | 70742581 | 64944333 |
| GH18_34 | 92815294 | 92432838 | 82966255 | 75637747 |
| GH18_35 | 112245909 | 111807969 | 96312275 | 86307334 |
| GH18_37 | 80292962 | 79989530 | 72075265 | 65773550 |
| GH18_38 | 89934948 | 89572941 | 80461283 | 73345462 |
| GH18_39 | 81837279 | 81511299 | 72327335 | 65789878 |
| GH18_40 | 91154587 | 90799797 | 80655947 | 73175635 |
| GH18_41 | 83616893 | 83288093 | 74338451 | 67466357 |
| GH18_42 | 92119863 | 91756620 | 80182515 | 72382085 |
| GH18_43 | 86487475 | 86230503 | 78664818 | 72096531 |
| GH18_45 | 68327717 | 68133761 | 62460905 | 57368692 |
| GH18_46 | 87349587 | 87040412 | 76251490 | 68859822 |
| GH18_49 | 103021745 | 102644042 | 92718288 | 84746617 |
| GH18_51 | 94632403 | 94256019 | 84343865 | 76837329 |
| GH18_52_01 | 63082765 | 62909800 | 57356059 | 52622841 |
| GH18_52_02 | 8153457 | 8130179 | 7396929 | 6787449 |
| GH18_53 | 69689268 | 69481517 | 63584413 | 58406581 |

| Sample ID | No. total reads | No. mapped reads | No. reads with mapping quality ≥ 20 | No. reads with mapping quality ≥ 60 |
|---|---|---|---|---|
| GH18_55 | 74391302 | 74167062 | 67446151 | 61875768 |
| GH18_59 | 66861130 | 66608707 | 59756899 | 54502858 |
| GH18_61 | 70713969 | 70493489 | 63812743 | 58420373 |
| GH18_62_01 | 61065851 | 60875735 | 55119498 | 50471054 |
| GH18_62_02 | 10712588 | 10680511 | 9632729 | 8820556 |
| GH18_66_01 | 56515693 | 56338129 | 50590251 | 46190131 |
| GH18_66_02 | 17922626 | 17863860 | 15994387 | 14604766 |
| GH18_67_01 | 64293330 | 64126123 | 58618260 | 53839373 |
| GH18_67_02 | 7825744 | 7804436 | 7117881 | 6538464 |
| GH18_68_01 | 64125923 | 63954795 | 58299055 | 53458230 |
| GH18_68_02 | 7588283 | 7566831 | 6884682 | 6315334 |
| GH18_69_01 | 57734839 | 57584602 | 52870362 | 48675456 |
| GH18_69_02 | 14565470 | 14529093 | 13298664 | 12245823 |
| GH18_72 | 76129214 | 75895557 | 68677524 | 62904260 |
| GH18_73 | 81252936 | 80981585 | 73424106 | 67287338 |
| GH18_74 | 76255517 | 76033299 | 69116707 | 63328771 |
| GH18_75 | 88361092 | 88082455 | 80366287 | 73736508 |
| GH18_76_01 | 63413352 | 63236583 | 57785005 | 53050946 |
| GH18_76_02 | 7227595 | 7207886 | 6565260 | 6028350 |
| GH18_77 | 66332675 | 66120240 | 60075746 | 55077984 |
| GH18_78 | 72591209 | 72340949 | 66063722 | 60712532 |
| IMC18_1_01 | 18617370 | 18569369 | 16998507 | 15645898 |
| IMC18_1_02 | 55278831 | 55110990 | 50452458 | 46375503 |
| IMC18_2_01 | 17070294 | 17023866 | 15538016 | 14280882 |
| IMC18_2_02 | 57290634 | 57111054 | 52153932 | 47862418 |
| IMC18_3_01 | 7126210 | 7104053 | 6489538 | 5970024 |
| IMC18_3_02 | 64220295 | 64009026 | 58519127 | 53804413 |
| IMC18_4_01 | 18490974 | 18429434 | 16830355 | 15466064 |
| IMC18_4_02 | 55625345 | 55420687 | 50642123 | 46488858 |
| IMC18_5_01 | 13666175 | 13609602 | 12398797 | 11380809 |
| IMC18_5_02 | 59592886 | 59333077 | 54059929 | 49557491 |
| IMC18_9 | 62618757 | 62256742 | 56073835 | 51381234 |
| IMC18_11 | 69134333 | 68925428 | 62865600 | 57720646 |
| IMC18_12 | 66757172 | 66548733 | 60779596 | 55830716 |
| IMC18_13 | 70557688 | 70309646 | 64734981 | 59664876 |
| IMC18_14 | 87507395 | 87181933 | 79734680 | 73258660 |
| IMC18_15_01 | 11060921 | 11027847 | 10089934 | 9289010 |
| IMC18_15_02 | 60095852 | 59904375 | 54877126 | 50488353 |
| IMC18_16_01 | 21952038 | 21881051 | 20035335 | 18455822 |
| IMC18_16_02 | 55602530 | 55402915 | 50754831 | 46692788 |
| IMC18_17_01 | 19909607 | 19835730 | 18136432 | 16676339 |
| IMC18_17_02 | 55951133 | 55732486 | 50973535 | 46805545 |
| IMC18_18_01 | 10037440 | 10006896 | 9163446 | 8430755 |

| Sample ID | No. total reads | No. mapped reads | No. reads with mapping quality ≥ 20 | No. reads with mapping quality ≥ 60 |
|---|---|---|---|---|
| IMC18_18_02 | 59099424 | 58892076 | 53746330 | 49314590 |
| IMC18_21 | 103323183 | 102928569 | 94050818 | 86418192 |
| IMC18_23_01 | 7436899 | 7416515 | 6798459 | 6256744 |
| IMC18_23_02 | 63290106 | 63098612 | 57920801 | 53283306 |
| IMC18_25 | 64179701 | 63927853 | 58289957 | 53548291 |
| IMC18_26_01 | 12556952 | 12519668 | 11411320 | 10496079 |
| IMC18_26_02 | 59555667 | 59356177 | 54126396 | 49715706 |
| IMC18_27_01 | 9351202 | 9321091 | 8504366 | 7811360 |
| IMC18_27_02 | 62139194 | 61925544 | 56590872 | 51952975 |
| IMC18_29_01 | 24093693 | 23959743 | 21898001 | 20127340 |
| IMC18_29_02 | 54907530 | 54584201 | 49910168 | 45802646 |
| IMC18_31_01 | 30919000 | 30832812 | 28188328 | 25920414 |
| IMC18_31_02 | 47701681 | 47558762 | 43495280 | 39948407 |
| IMC18_32_01 | 22483254 | 22424244 | 20454392 | 18806159 |
| IMC18_32_02 | 52452787 | 52296065 | 47734216 | 43826410 |
| IMC18_33 | 75278109 | 75031642 | 68583569 | 63022919 |
| IMC18_34 | 67385961 | 67137791 | 61547840 | 56672256 |
| IMC18_35 | 71051582 | 70755440 | 64362328 | 58958445 |
| IMC18_36 | 66179763 | 65952595 | 60460508 | 55568395 |
| IMC18_38 | 68378935 | 68095947 | 62448184 | 57434157 |
| IMC18_42 | 71203271 | 70954102 | 64678786 | 59281151 |
| IMC18_46_01 | 27827191 | 27736620 | 25230175 | 23174239 |
| IMC18_46_02 | 54515330 | 54298731 | 49491132 | 45423594 |
| MIL19_1_01 | 12873364 | 12834904 | 11678718 | 10717152 |
| MIL19_1_02 | 61118042 | 60942408 | 55682792 | 51088355 |
| MIL19_2_01 | 13098239 | 13059280 | 11924050 | 10968241 |
| MIL19_2_02 | 60909342 | 60734113 | 55656382 | 51199904 |
| MIL19_3 | 50669255 | 50437067 | 46051927 | 42313549 |
| MIL19_4 | 57325235 | 57071088 | 51870372 | 47523379 |
| MIL19_5_01 | 24028540 | 23927225 | 21940558 | 20230262 |
| MIL19_5_02 | 52070621 | 51873050 | 47627892 | 43918625 |
| MIL19_7 | 61372281 | 61122126 | 55684525 | 51025198 |
| MIL19_8_01 | 5295395 | 5271370 | 4814533 | 4420101 |
| MIL19_8_02 | 13989427 | 13919680 | 12738726 | 11683597 |
| MIL19_8_03 | 51067368 | 50832125 | 46364474 | 42600546 |
| MIL19_9_01 | 22593889 | 22522325 | 20461062 | 18783263 |
| MIL19_9_02 | 55709144 | 55542789 | 50673507 | 46524052 |
| MIL19_10_01 | 9890318 | 9853867 | 8961916 | 8184925 |
| MIL19_10_02 | 61723892 | 61511921 | 55682814 | 50941908 |
| MIL19_11_01 | 21520067 | 21460158 | 19540324 | 17941790 |
| MIL19_11_02 | 56200400 | 56052596 | 51217405 | 47027484 |
| MIL19_12_01 | 10621481 | 10578409 | 9651147 | 8815866 |
| MIL19_12_02 | 61200635 | 60983353 | 55366888 | 50696567 |

| Sample ID | No. total reads | No. mapped reads | No. reads with mapping quality ≥ 20 | No. reads with mapping quality ≥ 60 |
|---|---|---|---|---|
| MIL19_13 | 68473492 | 68253545 | 62133720 | 57045788 |
| MIL19_14 | 68078088 | 67823040 | 61593518 | 56436432 |
| MIL19_15 | 67964702 | 67724932 | 61547017 | 56408195 |
| MIL19_16 | 62896023 | 62644658 | 56933216 | 52168651 |
| MIL19_17_01 | 29996340 | 29908865 | 27298215 | 25075960 |
| MIL19_17_02 | 48253884 | 48127301 | 43960404 | 40387854 |
| MIL19_18_01 | 14492551 | 14451028 | 13119586 | 12029520 |
| MIL19_18_02 | 59646990 | 59484183 | 54231849 | 49729751 |
| MIL19_19 | 73849326 | 73565080 | 66887489 | 61339515 |
| MIL19_20_01 | 19751856 | 19661864 | 17977105 | 16587452 |
| MIL19_20_02 | 57162170 | 56916777 | 52324980 | 48274610 |
| MIL19_21 | 81044909 | 80744308 | 73157257 | 66922528 |
| MIL19_22_01 | 20191316 | 20115527 | 18475662 | 17064683 |
| MIL19_22_02 | 57404553 | 57204250 | 52615173 | 48595190 |
| MIL19_23_01 | 23094528 | 23005343 | 20792892 | 19063061 |
| MIL19_23_02 | 55723416 | 55547894 | 50456987 | 46304208 |
| MIL19_24 | 68299065 | 68053418 | 62122271 | 57053421 |
| MIL19_25_01 | 20747592 | 20688390 | 18868951 | 17340726 |
| MIL19_25_02 | 53331126 | 53183780 | 48718294 | 44771786 |
| MIL19_26 | 82987299 | 82711208 | 75283713 | 69023519 |
| MIL19_28 | 77925516 | 77687704 | 70921409 | 65189146 |
| MIL19_29_01 | 14942243 | 14898160 | 13587071 | 12487443 |
| MIL19_29_02 | 59902950 | 59732785 | 54633850 | 50230555 |
| MIL19_30_01 | 9428785 | 9386854 | 8554239 | 7871047 |
| MIL19_30_02 | 64322946 | 64041198 | 58629354 | 53938661 |
| MIL19_31 | 74152218 | 73878776 | 67215784 | 61655134 |
| MIL19_32_01 | 28214887 | 28108666 | 25622971 | 23601355 |
| MIL19_32_02 | 46867350 | 46710082 | 42816031 | 39452256 |
| MTC18_1_01 | 23564358 | 23492652 | 21547010 | 19855471 |
| MTC18_1_02 | 56185420 | 55984799 | 51453321 | 47378834 |
| MTC18_2_01 | 34982011 | 34854438 | 31875969 | 29351136 |
| MTC18_2_02 | 45752089 | 45560721 | 41777376 | 38438739 |
| MTC18_3_01 | 27227378 | 27144503 | 24955963 | 23022857 |
| MTC18_3_02 | 50936139 | 50763740 | 46762542 | 43118451 |
| MTC18_4_01 | 26007797 | 25912445 | 23624374 | 21716079 |
| MTC18_4_02 | 55741612 | 55493255 | 50698153 | 46561311 |
| MTC18_5 | 68309255 | 68025350 | 62529078 | 57754080 |
| MTC18_6_01 | 15774182 | 15725016 | 14454408 | 13348649 |
| MTC18_6_02 | 60489076 | 60271577 | 55512513 | 51219434 |
| MTC18_7 | 59213857 | 59004085 | 53756457 | 49222522 |
| MTC18_8_01 | 59458775 | 59264020 | 54233100 | 49889344 |
| MTC18_8_02 | 11961165 | 11917990 | 10893406 | 10016685 |
| MTC18_9 | 53711337 | 53257177 | 46997279 | 42341364 |

| Sample ID | No. total reads | No. mapped reads | No. reads with mapping quality ≥ 20 | No. reads with mapping quality ≥ 60 |
|---|---|---|---|---|
| MTC18_10 | 59239264 | 59033356 | 53802182 | 49249641 |
| MTC18_12_01 | 45435627 | 45292638 | 41239631 | 37810185 |
| MTC18_12_02 | 29468528 | 29380218 | 26668097 | 24442731 |
| MTC18_13 | 67837764 | 67066098 | 57589358 | 51949336 |
| MTC18_14 | 64186647 | 62724203 | 54042596 | 48441091 |
| MTC18_15_01 | 59052876 | 58851400 | 54183662 | 49961284 |
| MTC18_15_02 | 12238213 | 12191168 | 11205575 | 10326776 |
| MTC18_16 | 67564725 | 67280989 | 61429582 | 56422919 |
| MTC18_17 | 60970531 | 60749268 | 55558460 | 50956835 |
| MTC18_19 | 63651802 | 63366563 | 57203952 | 52365642 |
| MTC18_20 | 47257075 | 47043232 | 42736760 | 39172213 |
| MTC18_21 | 55107542 | 54880583 | 49674862 | 45484970 |
| MTC18_22 | 63642397 | 63246839 | 57069180 | 52363083 |
| MTC18_23 | 55470437 | 55305824 | 49297612 | 44650059 |
| MTC18_24 | 67598217 | 66592441 | 59474710 | 54080208 |
| MTC18_25 | 61631037 | 61127538 | 54180222 | 49152427 |
| MTC18_26_01 | 55203224 | 55046689 | 50418073 | 46329599 |
| MTC18_26_02 | 17306482 | 17250031 | 15775570 | 14489838 |
| MTC18_27_01 | 65989265 | 65651398 | 60762470 | 56162584 |
| MTC18_27_02 | 7590159 | 7548782 | 6978627 | 6447277 |
| MTC18_28_01 | 52698437 | 52532049 | 48325159 | 44537055 |
| MTC18_28_02 | 19880647 | 19808867 | 18179184 | 16741176 |
| MTC18_29 | 75709649 | 75002250 | 65719249 | 59273469 |
| MTC18_30 | 72520383 | 72290615 | 66651517 | 61449563 |
| MTC18_32 | 70361708 | 70035421 | 63568510 | 58141852 |
| MTC18_33 | 35006694 | 34865089 | 31650730 | 29016771 |
| MTC18_34_01 | 55905618 | 55715528 | 50987730 | 46773939 |
| MTC18_34_02 | 20574600 | 20506262 | 18715437 | 17159933 |
| NUB18_3 | 71895809 | 71574712 | 64958022 | 59497283 |
| NUB18_6 | 74480298 | 74178121 | 67585201 | 62018207 |
| NUB18_8 | 73647600 | 71092733 | 64088134 | 58586861 |
| NUB18_13 | 77617924 | 77321625 | 69950548 | 64053338 |
| NUB18_15 | 72498700 | 71493666 | 64783187 | 59393370 |
| NUB18_16 | 73883820 | 73651099 | 67115130 | 61564554 |
| NUB18_19 | 75499786 | 75081179 | 68212938 | 62619515 |
| NUB18_22 | 74760289 | 74541418 | 67240322 | 61464354 |
| NUB18_25 | 80064998 | 79826711 | 71624858 | 65541320 |
| NUB18_26 | 80820859 | 80490473 | 71577342 | 65325826 |
| NUB18_27 | 70265632 | 70020567 | 62481235 | 56991755 |
| NUB18_31 | 72461885 | 72174108 | 64192895 | 58496564 |
| NUB18_34 | 77546827 | 77303975 | 69488464 | 63502039 |
| NUB18_35 | 71882837 | 71636713 | 64447439 | 58984074 |
| NUB18_39 | 67939040 | 67739689 | 60708195 | 55472141 |

| Sample ID | No. total reads | No. mapped reads | No. reads with mapping quality ≥ 20 | No. reads with mapping quality ≥ 60 |
|---|---|---|---|---|
| NUB18_40 | 76747328 | 76543643 | 68787644 | 62932820 |
| NUB18_41 | 70443379 | 70191165 | 63260834 | 57949128 |
| NUB18_42_01 | 17345938 | 17255376 | 15541308 | 14227907 |
| NUB18_42_02 | 59589156 | 59242775 | 53468985 | 48924522 |
| NUB18_45_01 | 12547463 | 12498623 | 11387158 | 10473271 |
| NUB18_45_02 | 62494347 | 62219498 | 56809960 | 52222840 |
| NUB18_47_01 | 15359483 | 15317415 | 13794708 | 12627336 |
| NUB18_47_02 | 61227711 | 61022135 | 55054237 | 50353359 |
| NUB18_56 | 78847165 | 78314637 | 71205565 | 65308989 |
| NUB18_58_01 | 17942146 | 17868407 | 16125369 | 14774803 |
| NUB18_58_02 | 59292944 | 59008187 | 53359094 | 48843784 |
| NUB18_59_01 | 15426933 | 15374751 | 13945177 | 12804303 |
| NUB18_59_02 | 61033611 | 60793187 | 55239849 | 50695784 |
| NUB18_62_01 | 25849203 | 25782512 | 23422051 | 21483486 |
| NUB18_62_02 | 55108229 | 54941690 | 49981857 | 45816091 |
| NUB18_63_01 | 10698326 | 10664547 | 9598341 | 8776564 |
| NUB18_63_02 | 63940138 | 63697179 | 57434431 | 52459570 |
| NUB18_66_01 | 25604437 | 25509222 | 22654762 | 20586503 |
| NUB18_66_02 | 56026172 | 55778749 | 49678165 | 45111562 |
| NUB18_67_01 | 14582071 | 14447995 | 12983481 | 11859884 |
| NUB18_67_02 | 61463346 | 60851460 | 54789575 | 50005353 |
| NUB18_69_01 | 14452177 | 14386651 | 13026014 | 11955777 |
| NUB18_69_02 | 61517119 | 61202326 | 55535176 | 50934850 |
| NUB18_70_01 | 16474123 | 16425475 | 14840161 | 13595306 |
| NUB18_70_02 | 60568179 | 60357073 | 54654999 | 50041803 |
| NUB18_72_01 | 22827761 | 22736342 | 20453140 | 18704977 |
| NUB18_72_02 | 53467886 | 53224811 | 47977083 | 43824861 |

**Supplementary Table 4.** Numbers of single nucleotide polymorphisms (SNPs) out of Hardy-Weinberg equilibrium (HWE) in the within-group data set (MAF ≥ 0.05). A SNP is detected as out of HWE when its *P* value, calculated using VCFtools (0.1.16) (41), is smaller than the Bonferroni-corrected significance level at α=0.05 (*i.e.*, 0.05 divided by the total number of SNPs being tested).

| **Population** | **No. SNPs being tested** | **No. SNPs out of HWE** |
|---|---:|---:|
| Green Bay | 2,186,762 | 4,725 |
| Muskegon Lake | 2,275,931 | 4,548 |
| Grand Haven | 670,575 | 5,724 |
| Michigan City | 775,907 | 6,920 |
| Milwaukee | 776,082 | 6,186 |
| Naubinway | 711,223 | 5,877 |
| Sutton’s Bay | 727,014 | 3,725 |

**Supplementary Table 5.** Mean Weir and Cockerham's $F_{ST}$ between pairs of sample sites.

| | Green Bay | Muskegon Lake | Grand Haven | Michigan City | Milwaukee | Naubinway |
|---|---|---|---|---|---|---|
| Muskegon Lake | 0.051997 | | | | | |
| Grand Haven | 0.066478 | 0.058973 | | | | |
| Michigan City | 0.066747 | 0.059198 | 0.000195 | | | |
| Milwaukee | 0.063981 | 0.056034 | 0.000324 | -0.000186 | | |
| Naubinway | 0.059522 | 0.051000 | 0.000558 | 0.001302 | 0.000585 | |
| Sutton's Bay | 0.054967 | 0.046033 | 0.004133 | 0.003823 | 0.003257 | 0.001339 |

**Supplementary Table 6.** Sequencing depth of all samples in all sample sites.

| Population | Sample | Sequencing depth |
|---|---|---:|
| Green Bay | GBT1801 | 11.51 |
| Green Bay | GBT1802 | 11.97 |
| Green Bay | GBT1805 | 11.85 |
| Green Bay | GBT1806 | 11.74 |
| Green Bay | GBT1807 | 12.31 |
| Green Bay | GBT1808 | 14.25 |
| Green Bay | GBT1809 | 12.2 |
| Green Bay | GBT1810 | 12.08 |
| Green Bay | GBT1811 | 12.08 |
| Green Bay | GBT1812 | 11.85 |
| Green Bay | GBT1813 | 14.13 |
| Green Bay | GBT1814 | 11.74 |
| Green Bay | GBT1816 | 13.45 |
| Green Bay | GBT1818 | 13.11 |
| Green Bay | GBT1819 | 11.28 |
| Green Bay | GBT1820 | 14.59 |
| Green Bay | GBT1821 | 14.13 |
| Green Bay | GBT1824 | 14.82 |
| Green Bay | GBT1825 | 14.25 |
| Green Bay | GBT1826 | 13.68 |
| Green Bay | GBT1828 | 14.02 |
| Green Bay | GBT1829 | 16.64 |
| Green Bay | GBT1831 | 11.17 |
| Green Bay | GBT1832 | 13.68 |
| Green Bay | GBT1833 | 12.54 |
| Green Bay | GBT1834 | 11.97 |
| Green Bay | GBT1835 | 12.2 |
| Green Bay | GBU1809 | 13.45 |
| Green Bay | GBU1817 | 14.25 |
| Green Bay | GBU1821 | 13.11 |
| Muskegon Lake | D9320 | 11.63 |
| Muskegon Lake | D9321 | 12.54 |
| Muskegon Lake | D9322 | 11.28 |
| Muskegon Lake | D9323 | 12.2 |
| Muskegon Lake | D9324 | 11.97 |
| Muskegon Lake | D9325 | 11.51 |
| Muskegon Lake | D9326 | 12.42 |
| Muskegon Lake | D9327 | 13.45 |
| Muskegon Lake | D9328 | 12.99 |
| Muskegon Lake | D9329 | 11.4 |
| Muskegon Lake | D9330 | 12.2 |

| Population | Sample | Sequencing depth |
|---|---|---|
| Muskegon Lake | D9331 | 11.4 |
| Muskegon Lake | D9332 | 12.88 |
| Muskegon Lake | D9333 | 12.31 |
| Muskegon Lake | D9334 | 11.28 |
| Muskegon Lake | D9335 | 11.51 |
| Muskegon Lake | D9336 | 14.13 |
| Muskegon Lake | D9337 | 11.85 |
| Muskegon Lake | D9338 | 13.45 |
| Muskegon Lake | D9339 | 16.87 |
| Muskegon Lake | D9340 | 11.74 |
| Muskegon Lake | D9341 | 11.51 |
| Muskegon Lake | D9342 | 12.31 |
| Muskegon Lake | D9343 | 14.36 |
| Muskegon Lake | D9344 | 11.97 |
| Muskegon Lake | D9345 | 12.42 |
| Muskegon Lake | D9346 | 11.28 |
| Muskegon Lake | D9347 | 11.4 |
| Muskegon Lake | D9348 | 13.56 |
| Muskegon Lake | D9349 | 13.33 |
| Grand Haven | GH1834 | 15.61 |
| Grand Haven | GH1835 | 18.58 |
| Grand Haven | GH1837 | 13.45 |
| Grand Haven | GH1838 | 15.16 |
| Grand Haven | GH1839 | 13.68 |
| Grand Haven | GH1840 | 15.27 |
| Grand Haven | GH1841 | 14.02 |
| Grand Haven | GH1842 | 15.27 |
| Grand Haven | GH1843 | 14.59 |
| Grand Haven | GH1845 | 11.51 |
| Grand Haven | GH1846 | 14.47 |
| Grand Haven | GH1849 | 17.32 |
| Grand Haven | GH1851 | 15.84 |
| Grand Haven | GH1852 | 11.97 |
| Grand Haven | GH1853 | 11.74 |
| Grand Haven | GH1855 | 12.54 |
| Grand Haven | GH1859 | 11.28 |
| Grand Haven | GH1861 | 11.85 |
| Grand Haven | GH1862 | 12.08 |
| Grand Haven | GH1866 | 12.54 |
| Grand Haven | GH1867 | 12.2 |
| Grand Haven | GH1868 | 12.08 |
| Grand Haven | GH1869 | 12.2 |
| Grand Haven | GH1872 | 12.88 |

| Population | Sample | Sequencing depth |
|---|---|---|
| Grand Haven | GH1873 | 13.68 |
| Grand Haven | GH1874 | 12.88 |
| Grand Haven | GH1875 | 14.93 |
| Grand Haven | GH1876 | 11.97 |
| Grand Haven | GH1877 | 11.17 |
| Grand Haven | GH1878 | 12.31 |
| Michigan City | IMC1801 | 12.42 |
| Michigan City | IMC1802 | 12.54 |
| Michigan City | IMC1803 | 12.08 |
| Michigan City | IMC1804 | 12.54 |
| Michigan City | IMC1805 | 12.31 |
| Michigan City | IMC1809 | 10.49 |
| Michigan City | IMC1811 | 11.63 |
| Michigan City | IMC1812 | 11.28 |
| Michigan City | IMC1813 | 11.97 |
| Michigan City | IMC1814 | 14.7 |
| Michigan City | IMC1815 | 11.97 |
| Michigan City | IMC1816 | 13.11 |
| Michigan City | IMC1817 | 12.76 |
| Michigan City | IMC1818 | 11.63 |
| Michigan City | IMC1821 | 17.44 |
| Michigan City | IMC1823 | 11.97 |
| Michigan City | IMC1825 | 10.83 |
| Michigan City | IMC1826 | 12.2 |
| Michigan City | IMC1827 | 12.08 |
| Michigan City | IMC1829 | 13.33 |
| Michigan City | IMC1831 | 13.22 |
| Michigan City | IMC1832 | 12.65 |
| Michigan City | IMC1833 | 12.65 |
| Michigan City | IMC1834 | 11.4 |
| Michigan City | IMC1835 | 11.97 |
| Michigan City | IMC1836 | 11.17 |
| Michigan City | IMC1838 | 11.51 |
| Michigan City | IMC1842 | 12.08 |
| Michigan City | IMC1846 | 13.9 |
| Milwaukee | MIL1901 | 12.54 |
| Milwaukee | MIL1902 | 12.54 |
| Milwaukee | MIL1903 | 8.55 |
| Milwaukee | MIL1904 | 9.69 |
| Milwaukee | MIL1905 | 12.88 |
| Milwaukee | MIL1907 | 10.37 |
| Milwaukee | MIL1908 | 11.85 |
| Milwaukee | MIL1909 | 13.22 |

| Population | Sample | Sequencing depth |
|---|---|---|
| Milwaukee | MIL1910 | 12.08 |
| Milwaukee | MIL1911 | 13.11 |
| Milwaukee | MIL1912 | 12.08 |
| Milwaukee | MIL1913 | 11.51 |
| Milwaukee | MIL1914 | 11.51 |
| Milwaukee | MIL1915 | 11.4 |
| Milwaukee | MIL1916 | 10.6 |
| Milwaukee | MIL1917 | 13.22 |
| Milwaukee | MIL1918 | 12.54 |
| Milwaukee | MIL1919 | 12.42 |
| Milwaukee | MIL1920 | 12.99 |
| Milwaukee | MIL1921 | 13.68 |
| Milwaukee | MIL1922 | 13.11 |
| Milwaukee | MIL1923 | 13.22 |
| Milwaukee | MIL1924 | 11.51 |
| Milwaukee | MIL1925 | 12.54 |
| Milwaukee | MIL1926 | 14.02 |
| Milwaukee | MIL1928 | 13.11 |
| Milwaukee | MIL1929 | 12.65 |
| Milwaukee | MIL1930 | 12.42 |
| Milwaukee | MIL1931 | 12.54 |
| Milwaukee | MIL1932 | 12.65 |
| Naubinway | NUB1803 | 12.08 |
| Naubinway | NUB1806 | 12.54 |
| Naubinway | NUB1808 | 12.42 |
| Naubinway | NUB1813 | 13.11 |
| Naubinway | NUB1815 | 12.2 |
| Naubinway | NUB1816 | 12.42 |
| Naubinway | NUB1819 | 12.65 |
| Naubinway | NUB1822 | 12.54 |
| Naubinway | NUB1825 | 13.45 |
| Naubinway | NUB1826 | 13.56 |
| Naubinway | NUB1827 | 11.74 |
| Naubinway | NUB1831 | 12.2 |
| Naubinway | NUB1834 | 12.99 |
| Naubinway | NUB1835 | 12.08 |
| Naubinway | NUB1839 | 11.4 |
| Naubinway | NUB1840 | 12.88 |
| Naubinway | NUB1841 | 11.85 |
| Naubinway | NUB1842 | 12.99 |
| Naubinway | NUB1845 | 12.65 |
| Naubinway | NUB1847 | 12.88 |
| Naubinway | NUB1856 | 13.33 |

| Population | Sample | Sequencing depth |
|---|---|---|
| Naubinway | NUB1858 | 12.99 |
| Naubinway | NUB1859 | 12.88 |
| Naubinway | NUB1862 | 13.68 |
| Naubinway | NUB1863 | 12.54 |
| Naubinway | NUB1866 | 13.68 |
| Naubinway | NUB1867 | 12.76 |
| Naubinway | NUB1869 | 12.76 |
| Naubinway | NUB1870 | 12.99 |
| Naubinway | NUB1872 | 12.88 |
| Sutton's Bay | MTC1801 | 13.45 |
| Sutton's Bay | MTC1802 | 13.56 |
| Sutton's Bay | MTC1803 | 13.22 |
| Sutton's Bay | MTC1804 | 13.79 |
| Sutton's Bay | MTC1805 | 11.51 |
| Sutton's Bay | MTC1806 | 12.88 |
| Sutton's Bay | MTC1807 | 10.03 |
| Sutton's Bay | MTC1808 | 12.08 |
| Sutton's Bay | MTC1809 | 9 |
| Sutton's Bay | MTC1810 | 10.03 |
| Sutton's Bay | MTC1812 | 12.65 |
| Sutton's Bay | MTC1813 | 11.28 |
| Sutton's Bay | MTC1814 | 10.71 |
| Sutton's Bay | MTC1815 | 12.08 |
| Sutton's Bay | MTC1816 | 11.4 |
| Sutton's Bay | MTC1817 | 10.26 |
| Sutton's Bay | MTC1819 | 10.71 |
| Sutton's Bay | MTC1820 | 7.98 |
| Sutton's Bay | MTC1821 | 9.23 |
| Sutton's Bay | MTC1822 | 10.71 |
| Sutton's Bay | MTC1823 | 9.35 |
| Sutton's Bay | MTC1824 | 11.4 |
| Sutton's Bay | MTC1825 | 10.37 |
| Sutton's Bay | MTC1826 | 12.2 |
| Sutton's Bay | MTC1827 | 12.42 |
| Sutton's Bay | MTC1828 | 12.2 |
| Sutton's Bay | MTC1829 | 12.65 |
| Sutton's Bay | MTC1830 | 12.2 |
| Sutton's Bay | MTC1832 | 11.85 |
| Sutton's Bay | MTC1833 | 5.93 |
| Sutton's Bay | MTC1834 | 12.88 |

**Supplementary Table 7.** Mean read depth of all samples in all sample sites as calculated from the within-group data sets after alignment and genotyping (see “Variant calling and filtering” in Supplementary Materials for details).

| **Population** | **Sample** | **Mean read depth** |
|---|---|---:|
| Green Bay | GBT181 | 10.06 |
| Green Bay | GBT1810 | 9.99 |
| Green Bay | GBT1811 | 10.18 |
| Green Bay | GBT1812 | 9.52 |
| Green Bay | GBT1813 | 11.12 |
| Green Bay | GBT1814 | 9.38 |
| Green Bay | GBT1816 | 10.7 |
| Green Bay | GBT1818 | 10.43 |
| Green Bay | GBT1819 | 9.16 |
| Green Bay | GBT182 | 9.83 |
| Green Bay | GBT1820 | 11.67 |
| Green Bay | GBT1821 | 11.34 |
| Green Bay | GBT1824 | 12.16 |
| Green Bay | GBT1825 | 11.48 |
| Green Bay | GBT1826 | 11.18 |
| Green Bay | GBT1828 | 11.94 |
| Green Bay | GBT1829 | 13.12 |
| Green Bay | GBT1831 | 9.5 |
| Green Bay | GBT1832 | 11.17 |
| Green Bay | GBT1833 | 10.33 |
| Green Bay | GBT1834 | 9.72 |
| Green Bay | GBT1835 | 9.81 |
| Green Bay | GBT185 | 9.57 |
| Green Bay | GBT186 | 9.5 |
| Green Bay | GBT187 | 9.99 |
| Green Bay | GBT188 | 11.17 |
| Green Bay | GBT189 | 10.27 |
| Green Bay | GBU1817 | 12.3 |
| Green Bay | GBU1821 | 10.65 |
| Green Bay | GBU189 | 10.97 |
| Muskegon Lake | D9320 | 9.91 |
| Muskegon Lake | D9321 | 10.4 |
| Muskegon Lake | D9322 | 9.59 |
| Muskegon Lake | D9323 | 10.2 |
| Muskegon Lake | D9324 | 10.35 |
| Muskegon Lake | D9325 | 9.59 |
| Muskegon Lake | D9326 | 10.3 |
| Muskegon Lake | D9327 | 11.84 |
| Muskegon Lake | D9328 | 11.13 |

| Population | Sample | Mean read depth |
|---|---|---|
| Muskegon Lake | D9329 | 9.56 |
| Muskegon Lake | D9330 | 10.38 |
| Muskegon Lake | D9331 | 9.72 |
| Muskegon Lake | D9332 | 10.75 |
| Muskegon Lake | D9333 | 10.62 |
| Muskegon Lake | D9334 | 9.76 |
| Muskegon Lake | D9335 | 9.48 |
| Muskegon Lake | D9336 | 11.32 |
| Muskegon Lake | D9337 | 9.5 |
| Muskegon Lake | D9338 | 10.85 |
| Muskegon Lake | D9339 | 13.58 |
| Muskegon Lake | D9340 | 9.71 |
| Muskegon Lake | D9341 | 9.56 |
| Muskegon Lake | D9342 | 10.22 |
| Muskegon Lake | D9343 | 11.82 |
| Muskegon Lake | D9344 | 9.83 |
| Muskegon Lake | D9345 | 9.87 |
| Muskegon Lake | D9346 | 9.31 |
| Muskegon Lake | D9347 | 9.47 |
| Muskegon Lake | D9348 | 10.93 |
| Muskegon Lake | D9349 | 10.81 |
| Grand Haven | GH1834 | 14.76 |
| Grand Haven | GH1835 | 16.1 |
| Grand Haven | GH1837 | 13.25 |
| Grand Haven | GH1838 | 14.45 |
| Grand Haven | GH1839 | 12.84 |
| Grand Haven | GH1840 | 14.62 |
| Grand Haven | GH1841 | 13.54 |
| Grand Haven | GH1842 | 14.35 |
| Grand Haven | GH1843 | 13.37 |
| Grand Haven | GH1845 | 11.01 |
| Grand Haven | GH1846 | 13.12 |
| Grand Haven | GH1849 | 16.74 |
| Grand Haven | GH1851 | 15.34 |
| Grand Haven | GH1852 | 11.29 |
| Grand Haven | GH1853 | 11.31 |
| Grand Haven | GH1855 | 11.77 |
| Grand Haven | GH1859 | 10.92 |
| Grand Haven | GH1861 | 11.71 |
| Grand Haven | GH1862 | 11.69 |
| Grand Haven | GH1866 | 12.72 |
| Grand Haven | GH1867 | 11.78 |
| Grand Haven | GH1868 | 11.22 |

| Population | Sample | Mean read depth |
|---|---|---|
| Grand Haven | GH1869 | 11.69 |
| Grand Haven | GH1872 | 12.21 |
| Grand Haven | GH1873 | 12.98 |
| Grand Haven | GH1874 | 12.41 |
| Grand Haven | GH1875 | 13.88 |
| Grand Haven | GH1876 | 11.2 |
| Grand Haven | GH1877 | 10.87 |
| Grand Haven | GH1878 | 11.7 |
| Michigan City | IMC181 | 12.46 |
| Michigan City | IMC1811 | 11.06 |
| Michigan City | IMC1812 | 11.17 |
| Michigan City | IMC1813 | 11.58 |
| Michigan City | IMC1814 | 14.02 |
| Michigan City | IMC1815 | 11.78 |
| Michigan City | IMC1816 | 12.78 |
| Michigan City | IMC1817 | 12.85 |
| Michigan City | IMC1818 | 11.48 |
| Michigan City | IMC182 | 12.33 |
| Michigan City | IMC1821 | 15.9 |
| Michigan City | IMC1823 | 11.86 |
| Michigan City | IMC1825 | 10.11 |
| Michigan City | IMC1826 | 12.17 |
| Michigan City | IMC1827 | 11.76 |
| Michigan City | IMC1829 | 13.1 |
| Michigan City | IMC183 | 11.75 |
| Michigan City | IMC1831 | 13.18 |
| Michigan City | IMC1832 | 12.45 |
| Michigan City | IMC1833 | 11.85 |
| Michigan City | IMC1834 | 10.25 |
| Michigan City | IMC1835 | 11.26 |
| Michigan City | IMC1836 | 11.48 |
| Michigan City | IMC1838 | 11.61 |
| Michigan City | IMC184 | 12.24 |
| Michigan City | IMC1842 | 12.08 |
| Michigan City | IMC1846 | 14.42 |
| Michigan City | IMC185 | 11.95 |
| Michigan City | IMC189 | 9.97 |
| Milwaukee | MIL191 | 12.63 |
| Milwaukee | MIL1910 | 11.25 |
| Milwaukee | MIL1911 | 12.92 |
| Milwaukee | MIL1912 | 11.72 |
| Milwaukee | MIL1913 | 10.91 |
| Milwaukee | MIL1914 | 10.96 |

| Population | Sample | Mean read depth |
|---|---|---|
| Milwaukee | MIL1915 | 10.91 |
| Milwaukee | MIL1916 | 10.21 |
| Milwaukee | MIL1917 | 13.33 |
| Milwaukee | MIL1918 | 12.55 |
| Milwaukee | MIL1919 | 11.46 |
| Milwaukee | MIL192 | 12.38 |
| Milwaukee | MIL1920 | 13.06 |
| Milwaukee | MIL1921 | 13.08 |
| Milwaukee | MIL1922 | 13.4 |
| Milwaukee | MIL1923 | 13.23 |
| Milwaukee | MIL1924 | 10.97 |
| Milwaukee | MIL1925 | 12.41 |
| Milwaukee | MIL1926 | 13.24 |
| Milwaukee | MIL1928 | 12.39 |
| Milwaukee | MIL1929 | 12.56 |
| Milwaukee | MIL193 | 8.88 |
| Milwaukee | MIL1930 | 12.22 |
| Milwaukee | MIL1931 | 10.83 |
| Milwaukee | MIL1932 | 13.14 |
| Milwaukee | MIL194 | 8.5 |
| Milwaukee | MIL195 | 13.31 |
| Milwaukee | MIL197 | 9.64 |
| Milwaukee | MIL198 | 11.76 |
| Milwaukee | MIL199 | 12.94 |
| Naubinway | NUB1813 | 12.07 |
| Naubinway | NUB1815 | 11.39 |
| Naubinway | NUB1816 | 11.92 |
| Naubinway | NUB1819 | 12.26 |
| Naubinway | NUB1822 | 12.45 |
| Naubinway | NUB1825 | 12.73 |
| Naubinway | NUB1826 | 13.03 |
| Naubinway | NUB1827 | 11.42 |
| Naubinway | NUB183 | 11.41 |
| Naubinway | NUB1831 | 11.97 |
| Naubinway | NUB1834 | 12.71 |
| Naubinway | NUB1835 | 11.94 |
| Naubinway | NUB1839 | 11.4 |
| Naubinway | NUB1840 | 12.62 |
| Naubinway | NUB1841 | 12.06 |
| Naubinway | NUB1842 | 13.33 |
| Naubinway | NUB1845 | 13.15 |
| Naubinway | NUB1847 | 12.39 |
| Naubinway | NUB1856 | 12.44 |

| Population | Sample | Mean read depth |
|---|---|---|
| Naubinway | NUB1858 | 13.63 |
| Naubinway | NUB1859 | 13.2 |
| Naubinway | NUB186 | 11.8 |
| Naubinway | NUB1862 | 13.47 |
| Naubinway | NUB1863 | 13 |
| Naubinway | NUB1866 | 13.87 |
| Naubinway | NUB1867 | 13.07 |
| Naubinway | NUB1869 | 13.36 |
| Naubinway | NUB1870 | 13.39 |
| Naubinway | NUB1872 | 13.3 |
| Naubinway | NUB188 | 11.21 |
| Sutton’s Bay | MTC181 | 13.29 |
| Sutton’s Bay | MTC1810 | 9.5 |
| Sutton’s Bay | MTC1812 | 12.04 |
| Sutton’s Bay | MTC1813 | 9.42 |
| Sutton’s Bay | MTC1814 | 9.09 |
| Sutton’s Bay | MTC1815 | 10.94 |
| Sutton’s Bay | MTC1816 | 10.35 |
| Sutton’s Bay | MTC1817 | 7.17 |
| Sutton’s Bay | MTC1819 | 9.94 |
| Sutton’s Bay | MTC182 | 13.7 |
| Sutton’s Bay | MTC1820 | 7.9 |
| Sutton’s Bay | MTC1821 | 8.86 |
| Sutton’s Bay | MTC1822 | 9.6 |
| Sutton’s Bay | MTC1823 | 9.18 |
| Sutton’s Bay | MTC1824 | 9.95 |
| Sutton’s Bay | MTC1825 | 9.42 |
| Sutton’s Bay | MTC1826 | 11.41 |
| Sutton’s Bay | MTC1827 | 11.34 |
| Sutton’s Bay | MTC1828 | 11.67 |
| Sutton’s Bay | MTC1829 | 10.97 |
| Sutton’s Bay | MTC183 | 13.15 |
| Sutton’s Bay | MTC1830 | 11.29 |
| Sutton’s Bay | MTC1832 | 8.09 |
| Sutton’s Bay | MTC1833 | 6.04 |
| Sutton’s Bay | MTC1834 | 12.85 |
| Sutton’s Bay | MTC184 | 13.74 |
| Sutton’s Bay | MTC185 | 11.5 |
| Sutton’s Bay | MTC186 | 12.78 |
| Sutton’s Bay | MTC187 | 6.05 |
| Sutton’s Bay | MTC188 | 10.89 |
| Sutton’s Bay | MTC189 | 8.27 |

**Supplementary Table 8.** Contemporary estimates of *Ne* from GONE using whole-genome sequencing data.

| **Population** | ***Ne*** |
|---|---:|
| Green Bay | 46 |
| Muskegon Lake | 159 |
| Grand Haven | 4,173,580 |
| Michigan City | 4,532,970 |
| Milwaukee | 4,116,720 |
| Naubinway | 4,253,590 |
| Sutton’s Bay | 2,618,570 |

**Supplementary Table 9.** Genomic positions of, numbers of SNPs on, and numbers of SNPs with $F_{ST} \geq 0.6$ on 13 outlier windows.

| Outlier window | Chromosome | Start position | End position | No. SNPs | No. SNPs with $F_{ST} \geq 0.6$ |
|---|---|---|---|---|---|
| 1 | 1 | 2426937 | 2785691 | 3203 | 320 |
| 2 | 1 | 3544209 | 3739267 | 2290 | 163 |
| 3 | 1 | 15957452 | 16004538 | 363 | 141 |
| 4 | 4 | 34762415 | 34852528 | 2410 | 352 |
| 5 | 8 | 15887228 | 15918438 | 529 | 144 |
| 6 | 11 | 20466095 | 20615608 | 540 | 82 |
| 7 | 12 | 18929507 | 19043907 | 1173 | 184 |
| 8 | 13 | 21639613 | 21841952 | 1200 | 322 |
| 9 | 14 | 26911329 | 26930065 | 1265 | 194 |
| 10 | 18 | 14025882 | 14294333 | 1517 | 360 |
| 11 | 18 | 14911469 | 14941999 | 669 | 186 |
| 12 | 24 | 9892333 | 10102303 | 951 | 294 |
| 13 | 24 | 18658693 | 18978485 | 2449 | 190 |
